# Unique Amygdala Signatures and Shared Prefrontal Deficits in Autism: Mapping Social Heterogeneity via Naturalistic functional MRI

**DOI:** 10.64898/2026.02.26.708280

**Authors:** Xin Di, Ting Xu, Francisco Xavier Castellanos, Bharat B. Biswal

**Affiliations:** epartment of Biomedical Engineering, New Jersey Institute of Technology, Newark, NJ 07102, USA; Center for the Developing Brain, Child Mind Institute, New York, NY 10022, USA; Department of Child and Adolescent Psychiatry, NYU Grossman School of Medicine, New York, NY 10016, USA

**Keywords:** Autism spectrum disorder, amygdala, basal ganglia, medial prefrontal cortex, movie watching, principal component analysis

## Abstract

Naturalistic fMRI provides an ecologically valid window into social brain function, yet binary diagnostic labels may obscure neural signatures linked to the continuous spectrum of social deficits. We investigated whether social brain alterations in autism spectrum disorder (ASD) follow a categorical, dimensional, or “dual-track” architecture. We analyzed movie-watching fMRI data from the Healthy Brain Network cohort (ages 5–22) during two films: *The Present* (*n* = 327) and *Despicable Me* (*n* = 242), using principal component analysis (PCA) to quantify primary (PC1) and secondary (PC2) synchronization and variance partitioning to separate the contributions of categorical diagnosis from continuous symptom severity (Social Responsiveness Scale-2, SRS-2). During *The Present*, higher SRS-2 scores were associated with widespread reduction in synchronization. In social-motivational hubs (medial prefrontal cortex, caudate), reductions were carried by variance shared between diagnosis and SRS-2 scores, leaving the categorical and dimensional accounts empirically indistinguishable. The left amygdala was the only region with a unique dimensional association independent of diagnosis. Secondary response patterns (PC2) showed further unique dimensional associations in the precuneus, temporoparietal junction, and posterior superior temporal sulcus. No significant associations were detected during *Despicable Me*; a direct within-participant comparison (*n* = 198) localized reliable stimulus differences to the medial prefrontal and posterior cingulate cortex, with no difference in the amygdala. These findings identify dimension-specific signatures in the amygdala and in mentalizing-network timing, and indicate that the choice of stimulus affects whether such associations are detectable.

**Highlights:**

- Variance partitioning separates categorical from dimensional neural contributions.
- Medial prefrontal and caudate effects are shared by diagnosis and severity.
- Left amygdala uniquely tracks symptom severity beyond the diagnostic label.
- Secondary response patterns implicate mentalizing-network timing in autism.
- Prefrontal and posterior cingulate associations differ between two movie stimuli.

## 1. Introduction

Autism spectrum disorder (ASD) is a complex neurodevelopmental condition characterized by persistent difficulties in social communication and restricted, repetitive behaviors. While the prevalence of ASD has risen to approximately 1 in 36 children ^1^, the neural mechanisms underlying its profound phenotypic heterogeneity remain elusive. Traditional neuroimaging approaches, particularly resting-state functional MRI (fMRI), have successfully mapped intrinsic connectivity alterations in ASD ^2,3^. However, resting-state paradigms are inherently “blind” to the dynamic neural processing required for real-world social interaction. To truly capture the neural signatures of social impairment, it is necessary to probe the brain under conditions that simulate the complexity of daily life ^4^.

Naturalistic paradigms, such as movie watching, offer a powerful ecological alternative. Unlike static task-based fMRI, movies immerse participants in a continuous stream of social, emotional, and sensory information, acting as a “stress test” for the brain’s social processing circuitry ^5,6^. When typically developing (TD) individuals view the same engaging narrative, their brain activity exhibits highly synchronized patterns—referred to as inter-subject correlation (ISC)—in regions supporting sensory processing and mentalizing ^7^. In ASD, this synchronization is often disrupted, reflecting “idiosyncratic” processing of the shared world ^8^.

However, the landscape of naturalistic fMRI in ASD remains fragmented. While early studies reported reduced synchronization in the default mode network (DMN) and visual cortex ^8^, subsequent findings have been inconsistent, with some reporting widespread reductions ^9,10^ and others finding variable or even increased synchronization depending on the movie segment ^11,12^. These discrepancies likely stem from two critical limitations: small sample sizes (median *N* ≈ 20) and stimulus variability. The “social load” of the stimulus matters; cartoons with slapstick humor may not engage the same circuitry as emotionally complex narratives requiring theory of mind.

A more fundamental limitation of prior work is the reliance on binary case-control comparisons. ASD is inherently dimensional, with social communication deficits manifesting along a continuous spectrum ^13^. The Research Domain Criteria (RDoC) framework posits that neural circuits map more closely onto continuous behavioral dimensions than onto categorical diagnoses ^14^. Consequently, binary group comparisons may obscure critical “dual-track” relationships: some neural deficits may be strictly categorical (diagnosis-specific), while others—particularly in subcortical arousal systems—may track symptom severity (e.g., Social Responsiveness Scale [SRS] scores) regardless of the clinical label.

Furthermore, standard ISC analyses typically assume a single “canonical” response pattern (the group mean). This overlooks the possibility that individuals with ASD may exhibit structured heterogeneity—alternative modes of processing that are consistent across a subgroup but distinct from the typical response. Capturing these complex response patterns requires advanced decomposition methods, such as Principal Component Analysis (PCA), which can isolate the dominant shared signal (PC1) while simultaneously revealing secondary patterns (PC2) associated with temporal offset from the dominant response or idiosyncratic integration ^15^.

In the present study, we leveraged the Healthy Brain Network (HBN) dataset ^16^ to overcome these limitations. We analyzed fMRI data from children and adolescents watching two distinct animated films: *The Present* (Filmakademie Baden-Wuerttemberg, 2014), an emotionally complex narrative, and *Despicable Me* (Illumination, 2010), a slapstick comedy. At the primary head motion threshold, 327 participants contributed usable data for *The Present* (204 ASD, 123 TD) and 242 for *Despicable Me* (141 ASD, 101 TD), with 198 contributing both. This design allowed us to compare clinical associations across two stimuli. To ensure the robustness of our findings, we use a large sample size to detect subtle effects, strict head motion control with multiple sensitivity thresholds, and the parallel analysis of two different movie runs, compared directly within participants.

We employed a PCA-based variance partitioning framework to test two primary hypotheses. First, guided by the Social Motivation Theory ^17^ and the RDoC framework, we hypothesized a functional dissociation between cortical mentalizing regions and the amygdala. Prior work suggests that cortical regions supporting theory of mind (e.g., medial prefrontal cortex, mPFC) are consistently hypoconnected in ASD ^18^, reflecting the core cognitive deficits defined by the diagnostic category. In contrast, subcortical regions like the amygdala modulate social orienting and physiological arousal in a manner that often scales with symptom severity ^19,20^. Therefore, we predicted that while the mPFC would show variance shared between diagnosis and symptom severity, the amygdala would serve as a dimensional marker, tracking social impairment intensity independent of the categorical label.

Second, we hypothesized that secondary response patterns (PC2) would not be random noise but would capture meaningful variance related to the temporal integration of the narrative. We predicted that these secondary patterns would be particularly sensitive to dimensional severity, revealing differences in the timing of higher-order narrative integration.

## 2. Materials and Methods

### 2.1. Dataset and Participants

The MRI data were obtained from the Healthy Brain Network (HBN) project (http://fcon_1000.projects.nitrc.org/indi/cmi_healthy_brain_network/) ^16^. We identified 623 participants (Releases 1–11) with T1-weighted MRI scans, at least one run of movie-watching fMRI, and complete demographic and diagnostic information. To facilitate dimensional analyses, we included individuals with available Social Responsiveness Scale, Second Edition (SRS-2) scores. To harmonize data across age-specific forms (School-Age vs. Preschool), the SRS-2 Total T-score was used for all analyses.

Diagnostic classification was based on the consensus of clinicians using DSM-5 criteria. Participants were categorized into two groups: (1) ASD (confirmed diagnosis) and (2) TD (no psychiatric or neurological diagnoses). At the primary motion threshold, the sample for *The Present* comprised 327 participants (204 ASD, 123 TD) and for *Despicable Me* 242 (141 ASD, 101 TD); the lenient threshold retained 428 and 343 respectively (Table 1 and Supplementary Table S1).

**Table 1.**
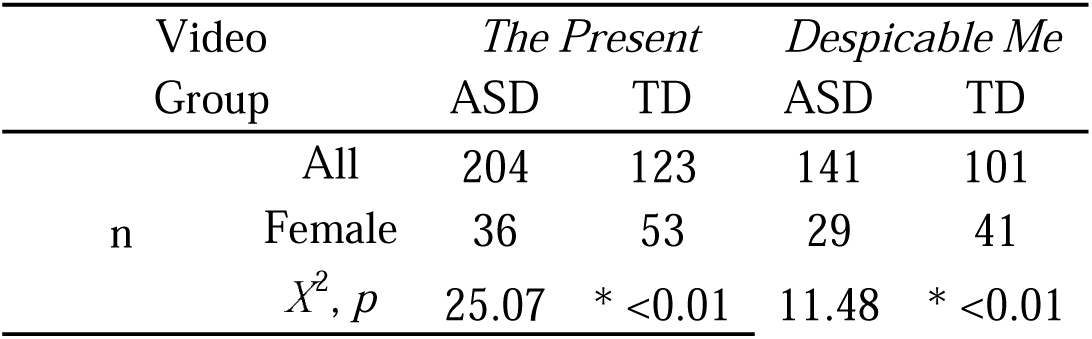
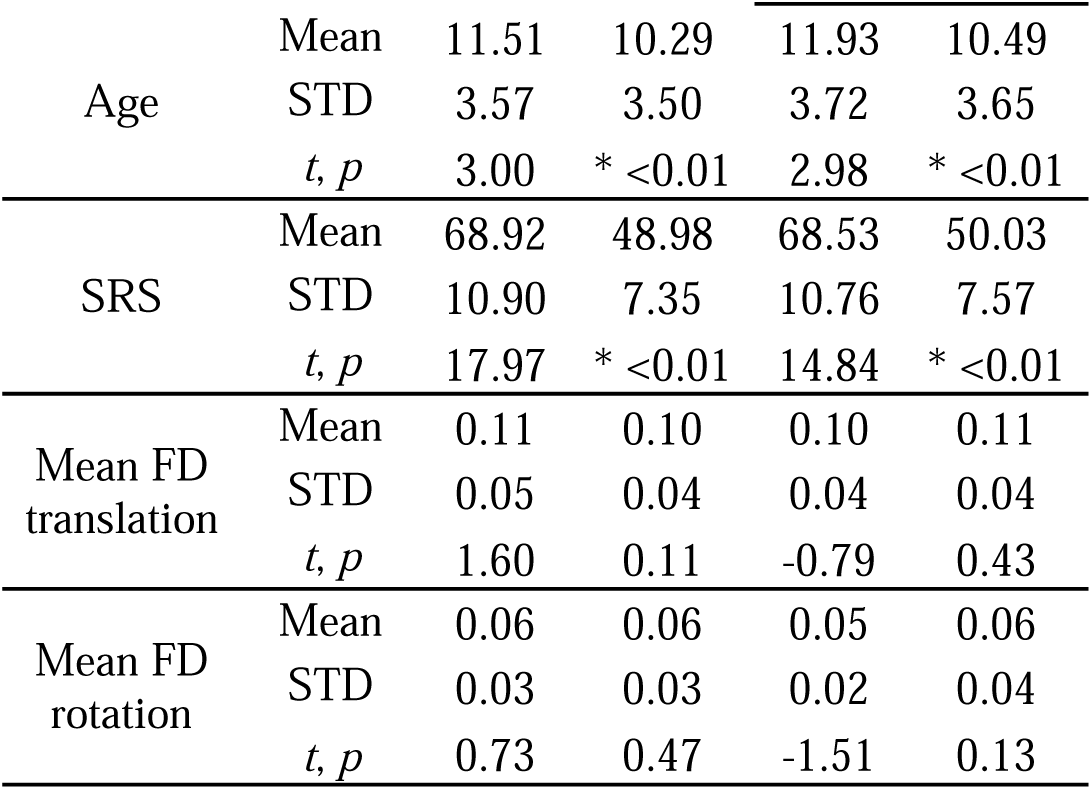
Participant characteristics at the primary head motion threshold (2.4 mm/°) for *The Present* and *Despicable Me*. Group comparisons used Χ² tests for sex and independent-samples *t* tests for age, Social Responsiveness Scale-2 (SRS-2) Total T-score, and mean framewise displacement (FD). * indicates *p* < 0.05 between the ASD and TD groups. Characteristics at the lenient (4.8 mm/°) and strict (1.2 mm/°) thresholds are provided in Supplementary Table S1.

Participants who completed both runs did so within the same scanning session. Because the longer *Despicable Me* run was more often incomplete or excluded for head motion, the two runs yielded overlapping but non-identical samples. At the primary motion threshold, 327 participants had usable data for *The Present*, 242 for *Despicable Me*, and 198 for both. Restricting both analyses to the overlapping subsample would have reduced the available samples by 39% and 18% respectively; we therefore analyzed each run in its maximal available sample and used the overlapping subsample only for the direct between-stimulus comparison, where a within-participant contrast is required. Participants contributing usable data for both runs did not differ from those contributing only *The Present* data in SRS-2 Total T-score, sex, or diagnosis, but were on average 1.5 years older (*p* < 0.001).

The original HBN data collection protocol was approved by the Chesapeake Institutional Review Board (now Advarra; protocol number: Pro00012309) and conducted in accordance with the Declaration of Helsinki. Written informed consent was obtained from all legal guardians, and written assent was obtained from all participants. The present study involved the secondary analysis of de-identified data and was determined to not require Institutional Review Board (IRB) review by the New Jersey Institute of Technology IRB (Protocol No. 2010003113) in accordance with U.S. Federal Regulations (45 CFR 46).

### 2.2. MRI Acquisition

Data were collected across three centers: Rutgers University Brain Imaging Center, (RUBIC), Citigroup Biomedical Imaging Center (CBIC), and City College of New York (CCNY), using 3T Siemens scanners (Trio and Prisma). Scanning protocols were harmonized across sites. Functional MRI was acquired using a multiband echo-planar imaging sequence: *TR* = 800 ms, *TE* = 30 ms, *flip angle* = 31°, *voxel size* = 2.4 x 2.4 x 2.4 mm^3^, and *multiband factor* = 6. This yielded 250 volumes for *The Present* (3:21 min) and 750 volumes for *Despicable Me* (10:00 min). High-resolution T1-weighted anatomical scans were acquired for registration.

### 2.3. Data Preprocessing

Preprocessing was performed using SPM12 (https://www.fil.ion.ucl.ac.uk/spm/) and standard MATLAB-based pipelines ^21^. T1-weighted images were segmented into tissue classes and normalized to Montreal Neurological Institute (MNI) space. For fMRI data, the first ten volumes were discarded to allow for signal stabilization. Remaining images were realigned, co-registered to the anatomical scan, normalized to MNI space (2.4 mm^3^ resolution), and smoothed with an 8 mm Gaussian kernel. Nuisance regression was applied using a General Linear Model (GLM) incorporating Friston’s 24-parameter motion model ^22^ and a high-pass filter (1/128 Hz). The resulting residual images were used for all subsequent analyses.

### 2.4. Motion Control and Sample Selection

To address the trade-off between data quality and sample representativeness in ASD, we adopted a multi-threshold strategy. We defined three maximum framewise displacement (FD) thresholds: lenient (4.8 mm/°), primary (2.4 mm/°), and strict (1.2 mm/°). The primary threshold (2.4 mm/°, equivalent to one voxel size) was used for main reporting, while the others served as sensitivity analyses to ensure robustness. In all group-level statistics, mean FD (translation and rotation) was included as a covariate.

### 2.5. Data analysis

#### 2.5.1. Parcellation

We analyzed 114 regions of interest (ROIs), combining the Schaefer 100-region cortical parcellation ^23^ with 14 bilateral subcortical regions from the Automated Anatomical Labeling (AAL) atlas ^24^, in AAL order and left before right within each pair: hippocampus (ROIs 101, 102), parahippocampal gyrus (103, 104), amygdala (105, 106), caudate (107, 108), putamen (109, 110), pallidum (111, 112), and thalamus (113, 114). For each preprocessed fMRI run, mean time series were extracted from all ROIs, yielding matrices of size 240 × 114 (*The Present*) or 740 × 114 (*Despicable Me*) for each participant.

#### 2.5.2. Analysis strategy across the two movie runs

Both runs were preprocessed with identical pipelines and submitted to the same decomposition and modelling framework. Neither run was designated as primary in advance, and the study was not pre-registered. Because the shared response is defined relative to a specific stimulus time course, PCA was performed separately for each run; components derived from different stimuli are not directly comparable, so the two runs were treated as parallel rather than pooled datasets. Results for both runs are reported in the main text using identical models and statistical criteria, including effect sizes for non-significant results.

We have previously analyzed these same two clips in parallel in this cohort, using overlapping but non-identical samples for the same reason, and found that individual differences in brain connectivity differed between the two clips in participants without psychiatric or neurological diagnoses ^25^. That observation motivated our interest in whether clinical associations would likewise differ between stimuli, which we test directly in Section 2.5.5.

#### 2.5.3. Inter-Subject Synchronization (PCA)

Shared neural responses were quantified using principal component analysis (PCA) ^15^. For each ROI, the time series of all *n* participants were assembled into a *t* × *n* matrix, and each participant’s time series (each column) was standardized (z-scored). The decomposition therefore operates on the correlation rather than the covariance structure, so that components reflect the consistency of response time courses across participants rather than differences in signal amplitude. PCA was performed along the time dimension to extract the dominant shared signal (PC1) and secondary patterns (PC2); the resulting loadings represent each participant’s synchronization with the group-level response.

PC1 loadings are closely related to conventional inter-subject correlation (ISC). In a previous methodological study using the same implementation, PC1 loadings were nearly identical to leave-one-out ISC values (*r* ≈ 1), and the correlation exceeded 0.95 whenever PC1 explained more than 5% of the variance ^15^. PC1 loadings can therefore be read as a close analogue of ISC, while the decomposition also yields higher-order components.

The variance explained by each component was tested against a null distribution generated by circular time-shift randomization ^15,26^. For each of 10,000 permutations, an independent shift was drawn for every participant from a discrete uniform distribution over [0, *T* − 1] with replacement (*T* = 240 for *The Present*, 740 for *Despicable Me*), each participant’s time series was circularly shifted by its own value, and PCA was reapplied as in the main analysis. This preserves the temporal autocorrelation of each individual time series while destroying the temporal correspondence between participants. The procedure established that the extracted components capture reliable shared responses (Figure 1); the group-level analyses of diagnosis and SRS-2 scores used the general linear models described below.

**Figure 1.**
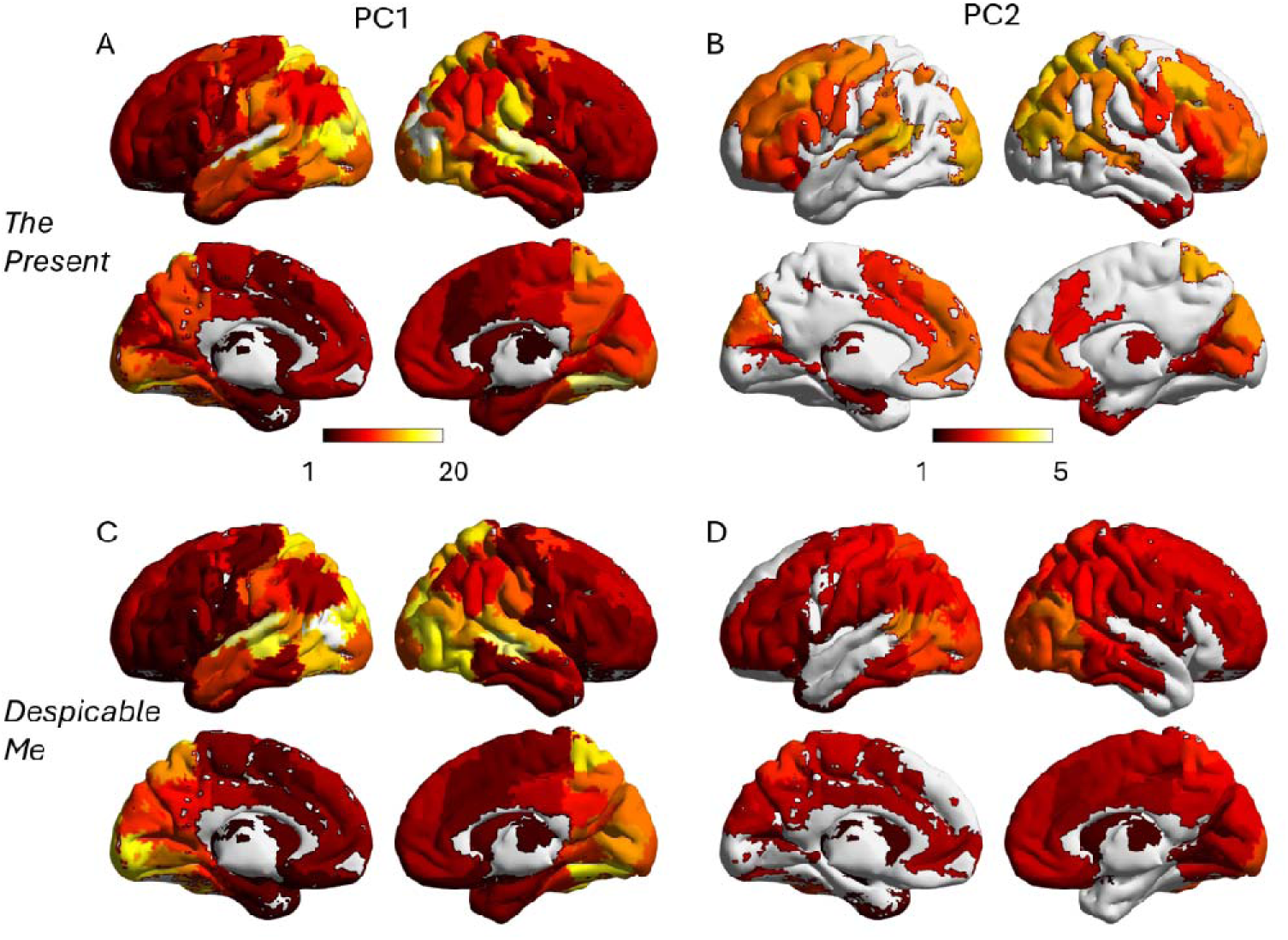
Spatial distribution of shared neural response patterns during *The Present* and *Despicable Me*. Maps show the percentage of variance explained by the primary (PC1; A, C) and secondary (PC2; B, D) principal components for *The Present* (A, B; *n* = 327) and *Despicable Me* (C, D; *n* = 242), at the primary head motion threshold of 2.4 mm/°. Note that the color scales differ between PC1 (1–20%) and PC2 (1–5%). Statistical significance was assessed using circular time-shift randomization with 10,000 permutations per region, thresholded at a false discovery rate (FDR) of *q* < 0.05; regions not reaching significance are shown in white. Cortical maps were visualized using BrainNet Viewer ^28^.

#### 2.5.4. General linear model

For each ROI, we performed general linear modeling (GLM) to examine the effects of categorical diagnosis and continuous autistic traits on regional synchronization, as indexed by the loadings of PC1 and PC2. To account for potential confounds, we first defined a set of standard covariates that were included in all analyses: age (modeled as log-age and log-age squared to account for non-linear developmental effects; ^15^), sex, scanning site, and log-transformed mean framewise displacement (head motion).

To systematically disentangle the effects of diagnosis and symptom severity, we constructed four distinct models using this same set of covariates:

1. Null Model: Covariates only
2. Diagnosis Model: Covariates + Diagnosis (binary).
3. SRS Model: Covariates + SRS-2 Total T-score (continuous).
4. Combined Model: Covariates + Diagnosis + SRS-2 Total T-score

The general structure of the full model is represented as:

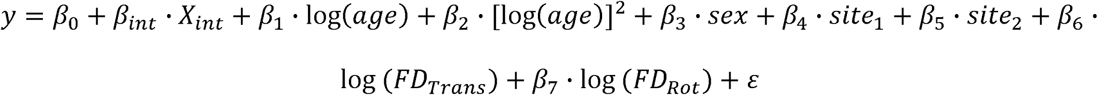

where *y* represents PC loadings of either the first or the second PC, and *X*_int_ represents the variable(s) of interest (Diagnosis, SRS-2, or both) depending on the model version. The models were fitted for each ROI, threshold, and movie clip.

To evaluate the impact of categorical diagnosis and continuous social traits, we examined the *t*-statistics associated with the β parameters for each variable of interest. The Combined Model was of primary interest, as it enabled the testing of unique associations with SRS-2 scores while strictly controlling for categorical diagnosis, and vice versa.

To formally quantify these relationships, we performed variance partitioning based on the coefficient of determination (*R*^2^) across the four nested models. We defined the Total Social Variance (*R*^2^_Total_) as the incremental explanatory power of both diagnosis and SRS-2 over the null model:

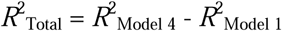

This metric represents the aggregate variance associated with social deficits after accounting for age, sex, site, and head motion. We then disentangled this total contribution into unique and shared components:

- Unique Diagnosis Variance: The variance explained by the diagnostic label that is not captured by SRS-2 scores (*R*^2^_Model 4_ - *R*^2^_Model 3_).
- Unique SRS Variance: The variance explained by SRS-2 scores that is not captured by the categorical diagnosis (*R*^2^_Model 4_ - *R*^2^_Model 2_).

Shared Variance: The overlapping information between diagnostic category and symptom severity, derived by subtracting the unique contributions from the total social variance:

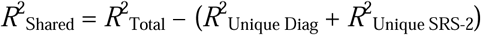

Due to the mathematical properties of *R*^2^ decomposition, the shared variance component can occasionally yield small negative values, typically indicative of suppression effects or negligible overlap between predictors.

Unique variance from the partitioning analysis is the squared semipartial correlation, a standard effect-size measure. Statistical inference proceeded in two stages. The SRS-only and diagnosis-only models were each tested across all 114 ROIs with FDR correction at *q* < 0.05. In the combined model, the unique contribution of each predictor was then evaluated only in the regions where that same predictor had reached significance in its single-predictor model — the unique SRS-2 effect in the SRS-2-significant regions, and the unique diagnosis effect in the diagnosis-significant regions. A single FDR threshold at *q* < 0.05 was computed across the pooled set of these retained tests. This nested procedure asks whether a marginally detectable association persists once the other predictor is controlled, rather than testing unique contributions in regions with no detectable association at all. Because the mask derives from the same data, the second stage constitutes conditional rather than independent inference, and effect estimates within it are subject to selection bias.

#### 2.5.5. Direct comparison between stimuli

To test whether clinical associations differed between stimuli, rather than inferring this from the presence versus absence of significance, we analyzed the 198 participants with usable data for both runs. For this analysis, PCA was recomputed within this subsample separately for each run, so that the group-level response for each stimulus was estimated from the same participants. Because loadings derived from separate PCAs are not on a common absolute scale, loadings were standardized across participants within each stimulus, and the within-participant difference (*The Present* − *Despicable Me*) was computed for each ROI. This difference was submitted to the same four nested models described above, with the between-run difference in mean framewise displacement as the motion covariate. Because age, sex, and site are constant across runs within a participant, these variables do not contribute to the within-participant contrast.

With both observations present for every participant, this difference-score model is algebraically equivalent to a mixed model including stimulus as a within-participant factor and a participant random intercept; the coefficient for SRS-2 in this model corresponds to the SRS-2 × stimulus interaction. We note that variance partitioning based on R² differences has no direct interaction analogue, so the difference-score model is a complementary framing rather than an extension of the same statistic.

Statistical correction proceeded hierarchically. We first applied FDR correction across all 114 ROIs. Where no region survived, we additionally restricted the correction to the 17 regions showing significant SRS-2 associations during *The Present* at the primary threshold. Because this restriction is derived from the same data, it constitutes a focused follow-up rather than independent confirmation, and effect estimates in these regions are subject to selection bias.

#### 2.5.6. Sensitivity and Developmental Analyses

To ensure the reliability and specificity of our findings, we performed five additional sets of analyses:

Head Motion Robustness: We verified that all primary findings (Diagnosis and SRS-2 effects) were consistent across the three maximum FD thresholds (1.2 mm/°, 2.4 mm/°, and 4.8 mm/°). This step ensured that the observed neural patterns were not artifacts of head motion or biased by the exclusion of more mobile participants.

Age-Matched Sensitivity Analysis: To rule out potential age-related confounds, we repeated the primary GLMs on a strictly age-balanced subsample (*n* = 123 for each group at 2.4 mm/° motion threshold). This subsample was generated using a greedy matching algorithm (see Supplementary Methods S1) that eliminated statistically significant age differences between the ASD and TD groups.

Mapping Developmental Trajectories: Beyond controlling for age as a covariate, we explicitly examined the independent contribution of age to brain synchronization. We calculated an F-contrast encompassing both the log(*age*) and [log(*age*)]^2^ parameters. This allowed us to map regions where synchronization varied significantly—either linearly or non-linearly—across the developmental span of 5–22 years. These developmental effects were examined for both PC1 and PC2 to test our hypothesis regarding the increased age-sensitivity of secondary response patterns.

Stimulus Duration: Because the two clips differ substantially in length, we divided the *Despicable Me* run into three consecutive segments of 240 volumes each, matching the duration of *The Present*, and repeated the primary analyses within each segment.

Age-Appropriate Tissue Priors: To evaluate whether findings depended on the use of adult-derived tissue probability maps, we regenerated priors matched to the age distribution of our sample using the Template-O-Matic toolbox^27^, which derives priors from the NIH MRI Study of Normal Brain Development, and repeated spatial normalization, ROI time series extraction, and the PCA-based analyses for both runs. A single sample-level prior was generated rather than participant-specific priors, as assigning different normalization targets by age would introduce age-dependent differences in deformation and effective ROI boundaries, confounding the age-related effects we model.

### 2.6. Design Considerations and Sample Size

Sample size was determined by data availability in the Healthy Brain Network cohort rather than by a priori power analysis (Section 2.1). For the primary cross-sectional regressions of a continuous predictor on regional synchronization, the available sample (*n* = 327) provides 80% power to detect a semipartial correlation of about 0.16 (≈2.5% unique variance) at α = 0.05, and about 0.20 (≈4%) under FDR correction across 114 regions — a range matched to the effects reported here, and considerably more sensitive than prior naturalistic fMRI studies of ASD (median *N* ≈ 20).

Rigor is further supported by design: both runs were analyzed under identical models; the between-stimulus comparison was performed within participants, holding age, sex, and site constant by construction at the cost of a smaller sample; and effects were evaluated at three motion thresholds and in an age-matched subsample. The study was not pre-registered, and estimates of unique variance are conditional on the regions selected for testing.

## 3. Results

### 3.1. Sample Characteristics

To control for head motion, we evaluated three maximum framewise displacement (FD) thresholds: 4.8, 2.4, and 1.2 mm/°. At the lenient threshold (4.8 mm/°), quality control retained 428 participants for *The Present* (262 ASD, 166 TD) and 343 for *Despicable Me* (202 ASD, 141 TD). Primary analyses used the 2.4 mm/° threshold, which retained 327 participants for *The Present* (204 ASD, 123 TD) and 242 for *Despicable Me* (141 ASD, 101 TD). Characteristics of these samples are given in Table 1 and Supplementary Figure S1. Characteristics at all three thresholds are provided in Supplementary Table S1.

In both samples, the ASD group included proportionally fewer females (*The Present*: *Χ*² = 25.07; *Despicable Me*: *Χ*² = 11.48; both *p* < 0.01) and was older than the TD group by approximately 1.2–1.4 years (*t* = 3.00 and 2.98, respectively; both *p* < 0.01). As expected, SRS-2 Total T-scores were substantially higher in the ASD group (*t* = 17.97 and 14.84; both *p* < 0.01). No group differences were observed in mean FD for either translation (*p* = 0.11 and 0.43) or rotation (*p* = 0.47 and 0.13), indicating no systematic motion bias. The same pattern held at the lenient and strict thresholds (Supplementary Table S1). Age, sex, scanning site, and mean FD were included as covariates in all subsequent analyses.

### 3.2. Shared Neural Synchronization

PC1 explained a significant proportion of variance in all 114 regions during both stimuli (FDR *q* < 0.05), with a similar spatial distribution across the two films (Figure 1A, C). The highest values occurred in visual, temporal, and parietal cortices, with lower contributions from prefrontal and subcortical regions. PC2 explained substantially less variance and reached significance in 59 regions during *The Present* and 93 regions during *Despicable Me* (Figure 1B, D). Although more regions reached significance during *Despicable Me*, the percentage of variance explained by PC2 was generally higher during *The Present*: the longer *Despicable Me* run affords greater sensitivity in the permutation test (740 versus 240 volumes), so the difference in the number of significant regions reflects statistical power rather than stronger secondary responses. PC2 was spatially distinct from PC1, involving bilateral frontoparietal regions, the mPFC, and the precuneus. Both components therefore captured reliable shared responses during both films, indicating that the absence of clinical associations during *Despicable Me* (Section 3.6) does not reflect a failure to recover a consistent group response to that stimulus.

### 3.3. Dimensional and Categorical Effects on Dominant Synchronization (PC1) during *The Present*

We examined the effects of continuous social traits (SRS-2 Total T-scores) and categorical diagnosis on PC1 loadings at the primary head motion threshold (2.4 mm/°).

#### Dimensional Analysis

Higher SRS-2 scores were associated with widespread reductions in neural synchronization (Figure 2A). At the primary threshold, significant negative associations were observed in 10 cortical and 7 subcortical regions. Twelve of these remained significant at all three motion thresholds (8 cortical, 4 subcortical; Figure 2B): bilateral orbitofrontal cortex (OFC; ROIs 31 and 79), left anterior temporal cortex (32 and 38), bilateral mPFC (44 and 96), bilateral posterior cingulate cortex (PCC; 50 and 100), left amygdala (105), bilateral caudate (107 and 108), and left putamen (109). The same regions were significant in the age-matched subsample (Figure S3A).

**Figure 2.**
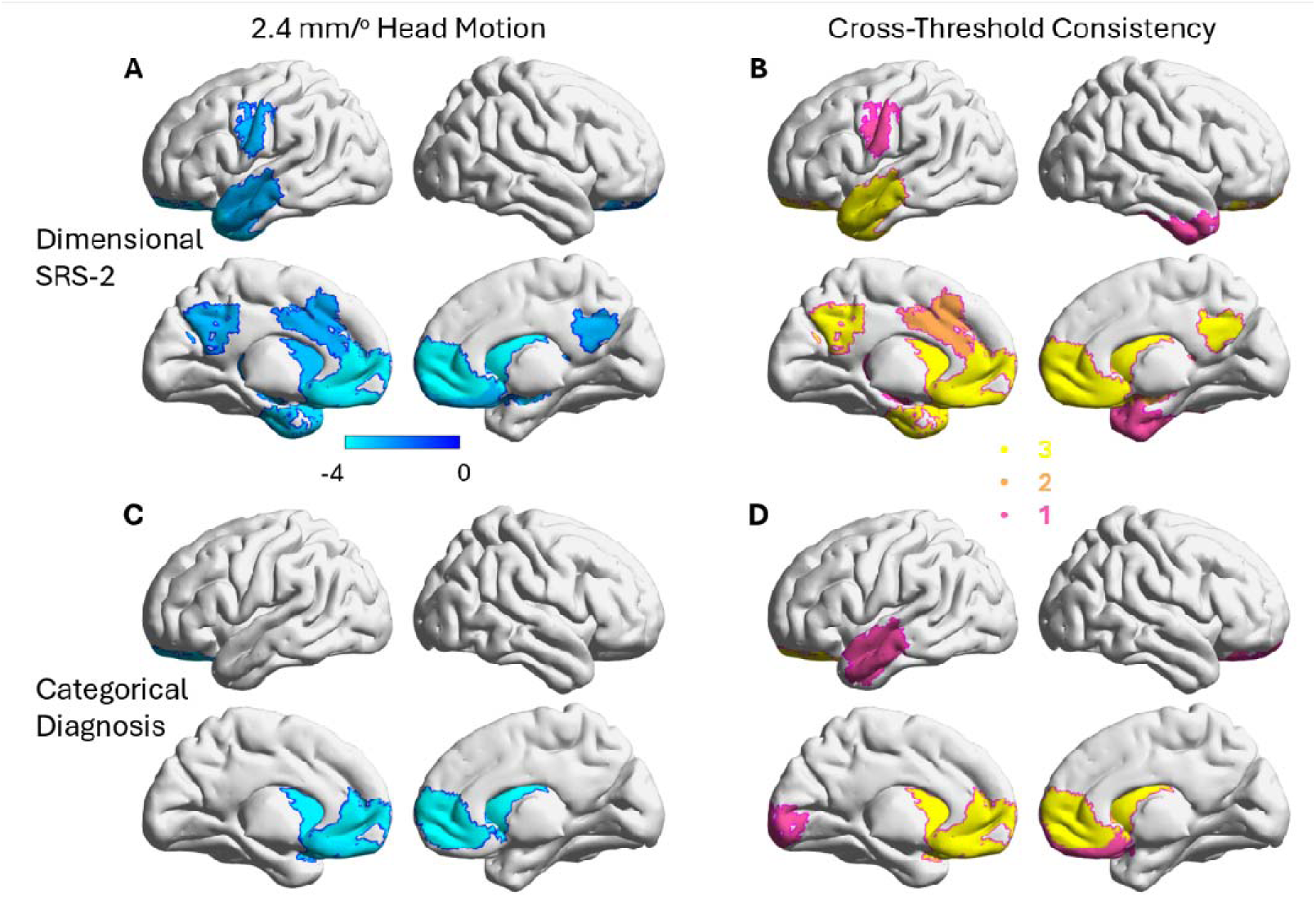
Comparison of dimensional and categorical effects on neural synchronization (PC1) during *The Present*. The top row illustrates the dimensional associations between Social Responsiveness Scale-2 (SRS-2) Total T-scores and regional synchronization (PC1 loadings), while the bottom row displays the categorical effects of clinical diagnosis (ASD vs. TD). Left panels present findings at the primary head motion threshold of 2.4 mm/°. Right panels show the consistency of these effects across three motion thresholds (1.2, 2.4, and 4.8 mm/°). The color scale represents the number of thresholds at which the effect remained significant, with values of 1, 2, and 3 corresponding to significance in one, two, or all three thresholds, respectively. All results were thresholded at a false discovery rate (FDR) of *q* < 0.05 across 114 regions of interest.

#### Categorical Analysis

Categorical comparison yielded a more restricted set of regions (Figure 2C). At the primary threshold, reduced synchronization in the ASD group was observed in five regions, which all remained significant at all three motion thresholds (Figure 2D): left OFC (31), bilateral mPFC (44 and 96), and bilateral caudate (107 and 108). The same regions were significant in the age-matched subsample (Figure S3B).

#### Variance Partitioning (Shared vs. Unique Effects)

In the combined model, the left amygdala (ROI 105) showed a significant unique association with SRS-2 scores independent of diagnosis (FDR *q* < 0.05 across the pooled masked tests), and was the only region, for either predictor, to reach significance at all three motion thresholds (Supplementary Figure S2). No region showed a significant unique contribution of categorical diagnosis across all three thresholds. Conversely, the diagnosis effects in the bilateral mPFC and caudate were largely explained by variance shared with SRS-2 scores. As visualized in Figure 3A, shared variance dominated the mPFC and caudate, whereas unique dimensional variance accounted for the majority of the effect in the amygdala. This effect was robust across all three motion thresholds (Supplementary Figure S2), in an age-matched analysis (Supplementary Section S2.2), and when spatial normalization was repeated using age-matched tissue probability maps (Supplementary Section S2.5).

**Figure 3.**
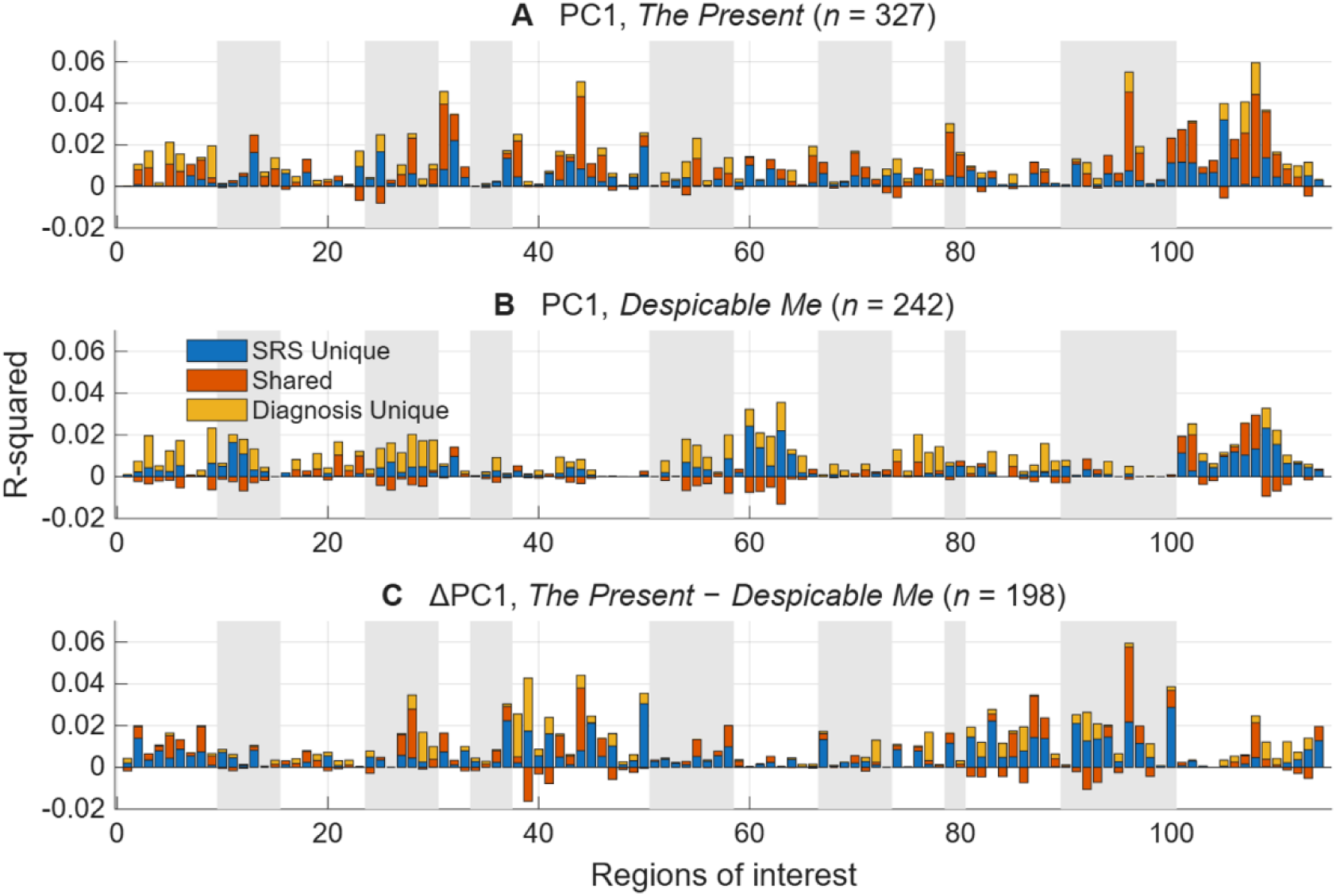
Variance partitioning of dimensional and categorical contributions to neural synchronization (PC1). Stacked bars show the decomposition of variance in PC1 loadings into the unique contribution of Social Responsiveness Scale-2 (SRS-2) Total T-scores, the unique contribution of clinical diagnosis (ASD vs. TD), and the variance shared between the two predictors, across 114 regions of interest. (A) *The Present*. (B) *Despicable Me*. (C) Within-participant difference in PC1 loadings (*The Present* − *Despicable Me*). Negative shared values indicate suppression, where the two predictors jointly explain more variance than the sum of their separate contributions. Data are shown for the primary head motion threshold (2.4 mm/°). Regions discussed in the text are identified by ROI index: mPFC (44, 96), PCC (50, 100), amygdala (105, 106), caudate (107, 108).

### 3.4. Dimensional and Categorical Effects on Dominant Synchronization (PC1) during *Despicable Me*

Applying the identical analytical framework to *Despicable Me* (*n* = 242), neither SRS-2 Total T-scores nor categorical diagnosis was significantly associated with PC1 loadings in any region at the primary motion threshold (FDR *q* < 0.05), and no region showed consistent effects across motion thresholds. Restricting the FDR correction to the 17 regions that showed significant SRS-2 associations during *The Present* at the primary threshold likewise yielded no significant effects.

Variance partitioning was consistent with these results (Figure 3B). Compared with *The Present*, the variance explained by dimensional, categorical, and combined models was substantially lower in bilateral default mode and limbic regions. Subcortical and visual regions showed a higher amount of variance compared with other regions (approximately 3%), although these effects were not significant.

We further split the whole data into three equal segments of 240 volumes to match with those in *The Present*. Again, neither SRS-2 Total T-scores nor categorical diagnosis was significantly associated with PC1 loadings in any region at the primary motion threshold in any of the three segments (FDR *q* < 0.05). Variance partitioning was not stable across segments: correlations between segments were close to zero for all three components, so we do not interpret the segment-level maps (Supplementary Section S2.3, Figure S4).

### 3.5. Direct comparison between stimuli

To test directly whether clinical associations differed between stimuli, rather than inferring this from the presence versus absence of significance, we analyzed the 198 participants with usable data for both stimuli. PC1 loadings were recomputed within this subsample for each stimulus and standardized within stimulus, and the within-participant difference (*The Present* − *Despicable Me*) was regressed on SRS-2 scores and diagnosis, with the difference in head motion between runs as a covariate. Because age, sex, and scanner are constant across runs, the contrast is unconfounded by these variables by construction.

No region survived FDR correction across all 114 regions. We therefore restricted the correction to the 17 regions showing significant SRS-2 associations during *The Present*; we note that this restriction is based on the same data and therefore constitutes a focused follow-up rather than independent confirmation. Within this set, three regions showed significant stimulus differences: right mPFC (ROI 96: *t* = −3.52, *p* = 0.0005), left mPFC (ROI 44: *t* = −2.81, *p* = 0.0054), and right PCC (ROI 100: *t* = −2.79, *p* = 0.0058). In each, the SRS-2 association was more negative during *The Present*. No region showed a significant stimulus difference for diagnosis.

By contrast, the left amygdala (ROI 105) showed no difference between stimuli (*t* = 0.25, *p* = 0.80), indicating that the apparent stimulus specificity of this association is not supported by direct comparison. Variance partitioning of the difference score was consistent with the regression results, indicating that these effects were carried predominantly by variance shared between SRS-2 scores and diagnosis (bilateral mPFC, ROI 44 and 96) or by the unique dimensional component (right PCC, ROI 100) (Figure 3C).

### 3.6. Dimensional and Categorical Effects on Secondary Neural Patterns (PC2)

Analysis of the secondary response pattern (PC2) revealed distinct dimensional associations. PC2 effects were generally less spatially extensive than PC1. The right posterior superior temporal sulcus (pSTS; ROI 93) showed a significant association with SRS-2 scores at all three motion thresholds, and the bilateral precuneus (ROIs 36 and 89) and left temporoparietal junction (TPJ; ROI 40) at two of the three (FDR *q* < 0.05 across all 114 regions; Figure 4). In the Schaefer parcellation, the precuneus ROIs are assigned to the control network and the temporoparietal and temporal ROIs to the default mode network. In the combined model, all four regions retained a significant unique association with SRS-2 scores after controlling for categorical diagnosis (FDR *q* < 0.05 across the pooled masked tests for both predictors). See Supplementary Figure S5 for the variance partition results at the primary threshold (2.4 mm/°) for PC2.

**Figure 4.**
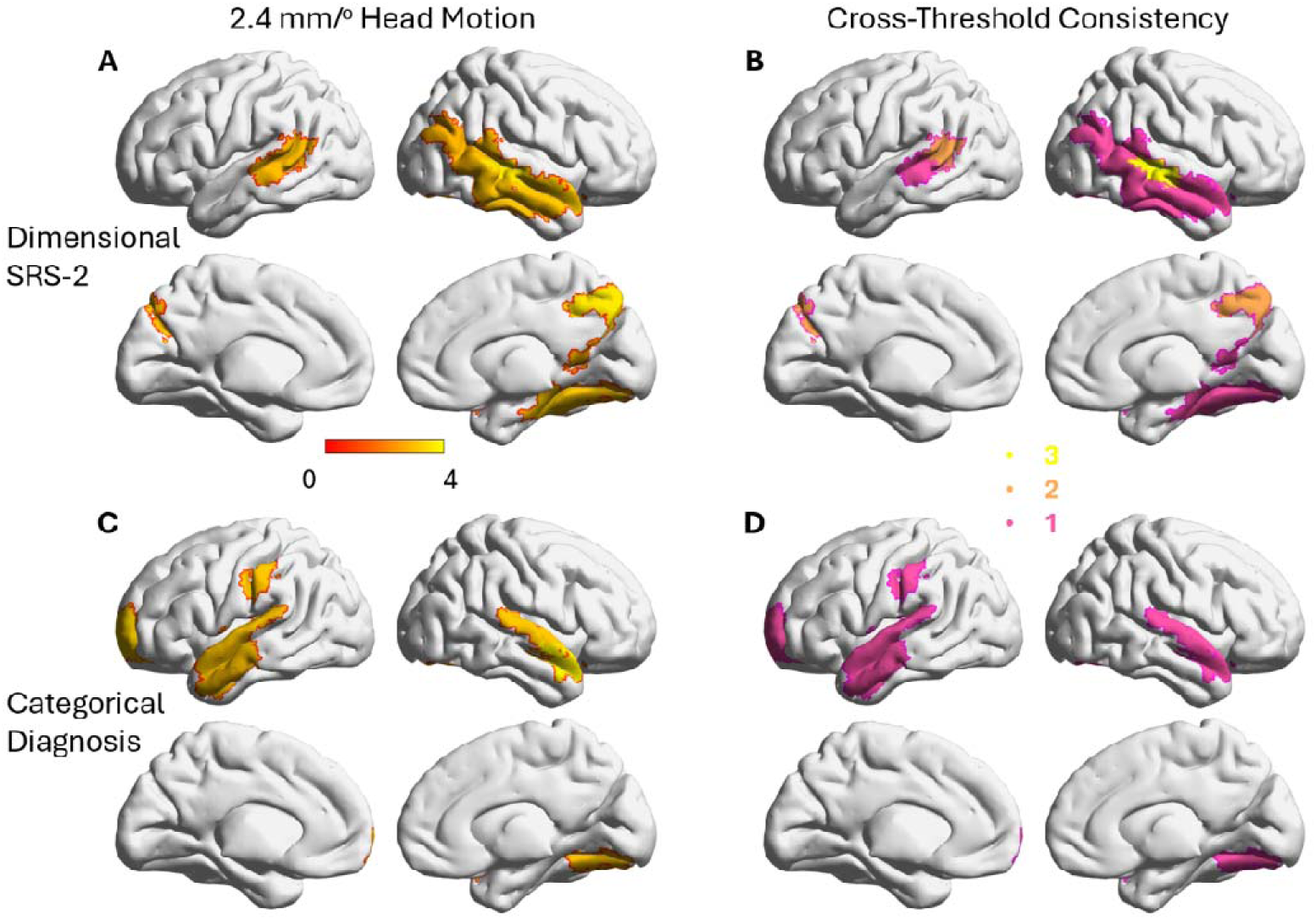
Comparison of dimensional and categorical effects on secondary shared response patterns (PC2) during *The Present*. The top row illustrates the dimensional associations between Social Responsiveness Scale-2 (SRS-2) Total T-scores and regional PC2 loadings, while the bottom row displays the categorical effects of clinical diagnosis (ASD vs. TD). Left panels present findings at the primary head motion threshold of 2.4 mm/°. Because the sign of PC2 loadings is mathematically arbitrary, the statistical maps represent absolute *t*-values. Right panels show the consistency of these effects across three motion thresholds (1.2, 2.4, and 4.8 mm/°). The color scale denotes the number of thresholds at which the effect remained significant, with values of 1, 2, and 3 corresponding to significance in one, two, or all three thresholds, respectively. All results were thresholded at a false discovery rate (FDR) of *q* < 0.05 across 114 regions of interest.

Categorical diagnosis was associated with PC2 loadings in seven regions at the primary threshold (ROIs 10, 17, 38, 45, 52, 59, and 92), but none reached significance at more than one threshold, and none retained a unique contribution after controlling for SRS-2 scores (Figure 4C, D).

Applying the same analysis to *Despicable Me*, neither SRS-2 scores nor diagnosis showed a significant association with PC2 loadings at the primary motion threshold (FDR *q* < 0.05 across 114 regions), and no region reached significance at more than one of the three thresholds. As with PC1, we therefore detected no dimensional or categorical associations with secondary responses during this stimulus. PC2 was not entered into the direct between-stimulus comparison, so these results do not establish a difference between stimuli.

### 3.7. Specificity of Clinical Effects vs. Developmental Maturation

To ensure that the identified signatures of social impairment were not confounded by the age differences between groups, we mapped the spatial distribution of age-related effects (Supplementary Figure S6). Age-related changes in PC1 synchronization were primarily localized to dorsal attention and somatomotor networks, showing no spatial overlap with the core social-motivational hubs (mPFC, caudate, and amygdala) associated with clinical measures. While PC2 patterns in the precuneus, TPJ, and pSTS showed sensitivity to both age and symptom severity, the reported SRS-2 effects represent unique variance explained after controlling for non-linear developmental trajectories. No significant main effects of sex were observed for either video stimulus.

## 4. Discussion

By applying variance partitioning to naturalistic fMRI data from a large developmental cohort (*n* = 327 during *The Present* and 242 during *Despicable Me*), we identified distinct neural signatures associated with the categorical diagnosis of ASD and the continuous dimension of social impairment. During *The Present,* our results demonstrate a functional dissociation: reduced synchronization in the mPFC and caudate reflects variance shared by both diagnostic labels and symptom severity, whereas the left amygdala serves as a unique dimensional marker, tracking social deficits independently of the categorical diagnosis. Secondary response patterns (PC2) in the precuneus, temporoparietal junction, and posterior superior temporal sulcus likewise showed a unique association with dimensional severity. Applying the same framework to *Despicable Me* yielded no significant associations in any region. A direct within-participant comparison of the two stimuli, however, localized reliable stimulus differences to the bilateral mPFC and right PCC only; the amygdala did not differ between stimuli, so the apparent stimulus specificity of the amygdala association is not supported by direct comparison.

### 4.1. The Amygdala as a Candidate Dimensional Marker

The left amygdala emerged as a unique dimensional hub. Unlike the mPFC and caudate, where diagnosis and SRS-2 scores captured overlapping information, the amygdala showed a significant unique association with SRS-2 scores after controlling for categorical diagnosis, and was the only region, for either predictor, whose unique contribution persisted across all three head motion thresholds. Amygdala synchronization therefore does not simply reflect the binary label. Because SRS-2 scores differed markedly between groups (Table 1), and the two predictors are correspondingly collinear, this unique component is estimated from variation in severity among participants sharing the same diagnostic status. It is this residual gradient, rather than a contrast between clinical and non-clinical individuals, that amygdala synchronization tracks.

While structural and functional alterations of the amygdala are established hallmarks of ASD ^29,30^, our variance partitioning refines this view: the amygdala’s relationship to social impairment conforms more closely to a dimensional than to a categorical model, consistent with the Research Domain Criteria framework ^14^. The direct between-stimulus comparison found no difference in the amygdala association between *The Present* and *Despicable Me*. Because that comparison used a smaller subsample (*n* = 198) and no significant association was detected during *Despicable Me* alone, we do not characterize the amygdala association as either stimulus-specific or stimulus-general; the present data do not distinguish these.

### 4.2. Convergence of Categorical and Dimensional Effects in the mPFC and Caudate

In contrast to the amygdala, the reductions in synchronization observed in the mPFC and caudate were predominantly carried by variance shared between diagnosis and SRS-2 scores. Neither predictor accounted for appreciable variance beyond the other, so these data do not distinguish a categorical from a dimensional account in these regions. This follows from how closely the two measures track one another in this sample (Table 1): where a region’s association is carried entirely by shared variance, the categorical and dimensional descriptions are empirically interchangeable, rather than one being more fundamental than the other ^13^.

These regions may therefore represent a “common core” of social deficits where categorical and dimensional models converge. The mPFC, a hub for self-referential thought and mentalizing, and the caudate, central to reward processing, are both implicated in the social motivation account of ASD ^17^, and their reduced synchronization is consistent with a generalized reduction in narrative tracking that varies with symptom severity. That alterations in both were observed within the same paradigm suggests naturalistic viewing engages social-cognitive and motivational processes concurrently, whereas task-based designs typically isolate them. The mPFC was also among the regions whose dimensional association differed between stimuli, which we consider next.

### 4.3. Secondary Response Patterns and Stimulus Dependency

PC2 effects were located in the bilateral precuneus (ROIs 36 and 89), assigned to the control network, and in the left TPJ (ROI 40) and right pSTS (ROI 93), both assigned to the default mode network. The latter two are canonical nodes of the mentalizing network, and together with the mPFC and PCC identified for PC1 they span much of the circuitry recruited when viewers attribute mental states during narrative. Because PC2 captures shared response patterns that are temporally offset from the dominant signal ^15^, a unique dimensional association in these regions suggests that social impairment involves not only weaker tracking of the shared narrative (PC1) but also altered timing of mentalizing-related integration. Because the sign of PC2 loadings is arbitrary, our data do not establish whether this reflects a lead or a lag. Flexible temporal modeling indicates that BOLD response profiles during naturalistic viewing are not well described by the canonical hemodynamic response ^31^, which makes a component indexing non-canonical temporal structure a plausible signal rather than residual noise.

For PC1, no significant associations were detected during *Despicable Me*. Because a null result cannot by itself establish a difference between stimuli, we compared the two directly in participants who viewed both. Dimensional associations were stronger during *The Present* in the bilateral mPFC and right PCC, but not in the amygdala. This regional selectivity is consistent with a “social stress test” account in which cortical mentalizing regions are most sensitive to the social-emotional load of the narrative: *The Present* requires empathy and theory of mind, whereas *Despicable Me* relies on slapstick humor and physical action. The two clips also differ in duration and in narrative completeness, however, so narrative complexity is one plausible contributor rather than an isolated factor, and synchronization is known to vary substantially across film stimuli ^32^. PC2 was not entered into the between-stimulus comparison, so we make no claim about its stimulus dependence.

### 4.4. Limitations

Several limitations warrant acknowledgment. First, our cross-sectional design limits inferences about developmental trajectories. Age effects on synchronization were spatially distinct from the clinical effects in the amygdala and mPFC (Supplementary Figure S6), which argues against the reported associations being a by-product of the group age difference, though it does not establish their mechanism. The age distribution is also right-skewed: 71% of participants were younger than 13 years and 2.8% older than 18, so the findings predominantly reflect childhood and early adolescence, and generalization to older adolescents and young adults cannot be established from these data.

Second, gaze behavior was not measured. Differences in where participants look can produce differences in synchronization independently of higher-order processing, and this applies to all regions we report. Future work integrating gaze-prediction models (e.g., CNN-based retrospective gaze estimation^33^) would help separate these contributions. Incorporating clinician-observed measures beyond caregiver reports (SRS-2) would likewise sharpen the dimensional characterization. Both the cortical and subcortical parcellations were derived from adults, and subcortical regions were taken from the atlas rather than individually segmented. No areal-level functional parcellation comparable to Schaefer-100 currently spans the 5–22 year range, and retaining these atlases preserves comparability with the adult literature against which our findings are interpreted. Agreement between standard and age-matched tissue priors was lowest in small structures adjacent to tissue boundaries, including the amygdala (Supplementary Section S2.5), so some uncertainty in the delineation of this region remains, although the reported association was unchanged under the age-matched pipeline.

Third, the effects we report are small in absolute terms (Figure 3). The size of the HBN sample is what made them detectable, consistent with large-scale analyses suggesting that reliable neural differences in ASD are subtle given the heterogeneity of the condition. We therefore emphasize the pattern of unique versus shared contributions rather than the magnitude of any single effect. Relatedly, because SRS-2 scores and diagnosis are strongly collinear in this sample, and because unique contributions were tested only in regions showing a marginal association, estimates of unique variance are conditional on that selection and describe the present sample rather than serving as unbiased effect sizes. A further selection concern is that the longer Despicable Me run and the motion thresholds themselves may preferentially exclude participants with more severe symptoms, which would attenuate dimensional associations. Participants contributing both runs did not differ from those contributing only The Present in SRS-2 score, sex, or diagnosis (Section 2.1), and the multi-threshold strategy was adopted partly for this reason (Supplementary Section S2.1), but residual selection of this kind cannot be excluded. These findings derive from a single cohort and have not been replicated in an independent sample; we therefore treat the left amygdala association as a candidate dimensional marker requiring replication rather than an established one.

Finally, medication information in HBN reflects substances taken on the day of the visit rather than chronic exposure, and was available for only 50 of 327 participants (15.3%); of these, 13 reported any medication and 5 a centrally acting agent, predominantly ADHD medications reflecting co-occurring conditions rather than treatment targeting core ASD features. We were therefore unable to evaluate whether psychotropic exposure influenced synchronization measures, and this possibility cannot be excluded.

## 5. Conclusion

By applying variance partitioning to large-scale naturalistic fMRI, this study identifies a functional dissociation within the social brain in autism spectrum disorder. We describe a “dual-track” architecture of social impairment: a shared social-motivational core in the mPFC and caudate, where categorical and dimensional descriptions are empirically interchangeable, and a unique dimensional signature in the left amygdala, which tracks graded variation in social-emotional impairment beyond what the diagnostic label captures. This dissociation does not follow the cortical–subcortical boundary: the caudate patterns with the mPFC rather than with the amygdala. These findings are consistent with the RDoC framework, in which continuous behavioral dimensions may map onto neural circuits more closely than diagnostic categories do, and they suggest that whether a categorical or a dimensional account fits better is a question to be settled region by region.

Secondary response patterns (PC2) revealed a further dimension-specific association in the precuneus, temporoparietal junction, and posterior superior temporal sulcus, nodes of the mentalizing network, indicating that social impairment involves altered timing of higher-order narrative integration as well as weaker tracking of the shared response. Dimensional associations in prefrontal and cingulate cortex were stronger during *The Present* than during *Despicable Me*, which suggests that the choice of stimulus materially affects whether such associations are detectable at all. Combining ecologically valid paradigms with dimensional analytics therefore offers a route toward quantitative neurobiological markers of social impairment, provided the stimuli themselves are selected and characterized as carefully as the measures.

## Acknowledgements

We acknowledge the computing resources provided by the high-performance computing (HPC) facility at NJIT.

## Author Contributions

**Xin Di:** Conceptualization, Methodology, Software, Formal analysis, Visualization, Project administration, Funding acquisition, Writing – original draft, Writing – review & editing.

**Ting Xu:** Resources, Writing – review & editing.

**Francisco Xavier Castellanos**: Writing – review & editing.

**Bharat B. Biswal:** Resources, Supervision, Funding acquisition, Writing – review & editing.

## Declaration of generative AI and AI-assisted technologies in the manuscript preparation process

During the preparation of this work, the authors used Claude, Gemini, and ChatGPT to improve the readability, grammar, and flow of the manuscript text. After using this tool, the authors reviewed and edited the content as needed and take full responsibility for the content of the publication.

## Reference

1. Maenner MJ. Prevalence and Characteristics of Autism Spectrum Disorder Among Children Aged 8 Years — Autism and Developmental Disabilities Monitoring Network, 11 Sites, United States, 2020. MMWR Surveill Summ. 2023;72. doi:10.15585/mmwr.ss7202a1

2. Di Martino a, Yan CG, Li Q, et al. The autism brain imaging data exchange: towards a large-scale evaluation of the intrinsic brain architecture in autism. Molecular psychiatry. 2014;19(April):659–667. doi:10.1038/mp.2013.78

3. Hull JV, Dokovna LB, Jacokes ZJ, Torgerson CM, Irimia A, Van Horn JD. Resting-State Functional Connectivity in Autism Spectrum Disorders: A Review. Front Psychiatry. 2017;7. doi:10.3389/fpsyt.2016.00205

4. Eickhoff SB, Milham M, Vanderwal T. Towards clinical applications of movie fMRI. NeuroImage. 2020;217:116860. doi:10.1016/j.neuroimage.2020.116860

5. Finn ES, Bandettini PA. Movie-watching outperforms rest for functional connectivity-based prediction of behavior. NeuroImage. 2021;235:117963. doi:10.1016/j.neuroimage.2021.117963

6. Richardson H, Lisandrelli G, Riobueno-Naylor A, Saxe R. Development of the social brain from age three to twelve years. Nat Commun. 2018;9(1):1–12. doi:10.1038/s41467-018-03399-2

7. Hasson U, Nir Y, Levy I, Fuhrmann G, Malach R. Intersubject synchronization of cortical activity during natural vision. Science (New York, NY). 2004;303(5664):1634–1640. doi:10.1126/science.1089506

8. Hasson U, Avidan G, Gelbard H, et al. Shared and idiosyncratic cortical activation patterns in autism revealed under continuous real-life viewing conditions. Autism Research. 2009;2(4):220–231. doi:10.1002/aur.89

9. Byrge L, Dubois J, Tyszka JM, Adolphs R, Kennedy DP. Idiosyncratic Brain Activation Patterns Are Associated with Poor Social Comprehension in Autism. J Neurosci. 2015;35(14):5837–5850. doi:10.1523/JNEUROSCI.5182-14.2015

10. Lyons KM, Stevenson RA, Owen AM, Stojanoski B. Examining the relationship between measures of autistic traits and neural synchrony during movies in children with and without autism. NeuroImage: Clinical. 2020;28:102477. doi:10.1016/j.nicl.2020.102477

11. Mangnus M, Koch SBJ, Cai K, et al. Preserved Spontaneous Mentalizing Amid Reduced Intersubject Variability in Autism During a Movie Narrative. Biological Psychiatry: Cognitive Neuroscience and Neuroimaging. Published online October 28, 2024. doi:10.1016/j.bpsc.2024.10.007

12. Ou W, Zeng W, Gao W, et al. Movie Events Detecting Reveals Inter-Subject Synchrony Difference of Functional Brain Activity in Autism Spectrum Disorder. Front Comput Neurosci. 2022;16. doi:10.3389/fncom.2022.877204

13. Constantino JN, Todd RD. Autistic Traits in the General Population: A Twin Study. Arch Gen Psychiatry. 2003;60(5):524–530. doi:10.1001/archpsyc.60.5.524

14. Cuthbert BN, Insel TR. Toward the future of psychiatric diagnosis: the seven pillars of RDoC. BMC Med. 2013;11(1):126. doi:10.1186/1741-7015-11-126

15. Di X, Biswal BB. Principal component analysis reveals multiple consistent responses to naturalistic stimuli in children and adults. Human Brain Mapping. 2022;43(11):3332–3345. doi:10.1002/hbm.25568

16. Alexander LM, Escalera J, Ai L, et al. An open resource for transdiagnostic research in pediatric mental health and learning disorders. Scientific Data. 2017;4(1):1–26. doi:10.1038/sdata.2017.181

17. Chevallier C, Kohls G, Troiani V, Brodkin ES, Schultz RT. The social motivation theory of autism. Trends in Cognitive Sciences. 2012;16(4):231–239. doi:10.1016/j.tics.2012.02.007

18. Kennedy DP, Courchesne E. The intrinsic functional organization of the brain is altered in autism. NeuroImage. 2008;39(4):1877–1885. doi:10.1016/j.neuroimage.2007.10.052

19. Dalton KM, Nacewicz BM, Johnstone T, et al. Gaze fixation and the neural circuitry of face processing in autism. Nat Neurosci. 2005;8(4):519–526. doi:10.1038/nn1421

20. Kleinhans NM, Johnson LC, Richards T, et al. Reduced Neural Habituation in the Amygdala and Social Impairments in Autism Spectrum Disorders. American Journal of Psychiatry. 2009;166:467–475. doi:10.1176/appi.ajp.2008.07101681

21. Di X, Biswal BB. A functional MRI pre-processing and quality control protocol based on statistical parametric mapping (SPM) and MATLAB. Frontiers in Neuroimaging. 2023;1:1070151.

22. Friston KJ, Williams S, Howard R, Frackowiak RS, Turner R. Movement-related effects in fMRI time-series. Magnetic resonance in medicine-: official journal of the Society of Magnetic Resonance in Medicine / Society of Magnetic Resonance in Medicine. 1996;35(3):346–355. doi:DOI 10.1002/mrm.1910350312

23. Schaefer A, Kong R, Gordon EM, et al. Local-Global Parcellation of the Human Cerebral Cortex from Intrinsic Functional Connectivity MRI. Cerebral cortex (New York, NY-: 1991). 2018;28(9):3095-3114. doi:10.1093/cercor/bhx179

24. Tzourio-Mazoyer N, Landeau B, Papathanassiou D, et al. Automated anatomical labeling of activations in SPM using a macroscopic anatomical parcellation of the MNI MRI single-subject brain. NeuroImage. 2002;15(1):273–289. doi:10.1006/nimg.2001.0978

25. Di X, Xu T, Uddin LQ, Biswal BB. Individual differences in time-varying and stationary brain connectivity during movie watching from childhood to early adulthood: Age, sex, and behavioral associations. Developmental Cognitive Neuroscience. 2023;63:101280. doi:10.1016/j.dcn.2023.101280

26. Kauppi JP, Jääskeläinen IP, Sams M, Tohka J. Inter-subject correlation of brain hemodynamic responses during watching a movie: localization in space and frequency. Front Neuroinform. 2010;4. doi:10.3389/fninf.2010.00005

27. Wilke M, Holland SK, Altaye M, Gaser C. Template-O-Matic: a toolbox for creating customized pediatric templates. Neuroimage. 2008;41(3):903–913. doi:10.1016/j.neuroimage.2008.02.056

28. Xia M, Wang J, He Y. BrainNet Viewer: a network visualization tool for human brain connectomics. Csermely P, ed. PloS one. 2013;8(7):e68910. doi:10.1371/journal.pone.0068910

29. Baron-Cohen S, Ring HA, Bullmore ET, Wheelwright S, Ashwin C, Williams SCR. The amygdala theory of autism. Neuroscience & Biobehavioral Reviews. 2000;24(3):355–364. doi:10.1016/S0149-7634(00)00011-7

30. Pelphrey KA, Shultz S, Hudac CM, Vander Wyk BC. Research Review: Constraining heterogeneity: the social brain and its development in autism spectrum disorder. Journal of Child Psychology and Psychiatry. 2011;52(6):631–644. doi:10.1111/j.1469-7610.2010.02349.x

31. Di X, Hanna GB, Biswal BB. Beyond the Canonical HRF: Flexible Temporal Modeling Reveals Unconstrained BOLD Profiles During Naturalistic Viewing. Imaging Neuroscience. Published online July 17, 2026. doi:10.1162/IMAG.a.1331

32. Leipold S, Rao RR, Schoffelen JM, Bögels S, Toni I. Inter-subject correlations and their behavioral associations vary across movies: Implications for generalizability. Imaging Neuroscience. Published online September 25, 2026. doi:10.1162/IMAG.a.1400

33. Gao L, Wei Z, Biswal BB, Di X. fMRI-based prediction of eye gaze during naturalistic movie viewing reveals eye-movement-related brain activity. psychoradiology. 2026;6:kkag026. doi:10.1093/psyrad/kkag026

34. Lombardo MV, Lai MC, Baron-Cohen S. Big data approaches to decomposing heterogeneity across the autism spectrum. Mol Psychiatry. 2019;24(10):1435–1450. doi:10.1038/s41380-018-0321-0

