## Supplementary Materials for "Unique Amygdala Signatures and Shared Prefrontal Deficits in Autism: Mapping Social Heterogeneity via Naturalistic functional MRI"

**Table of Contents**

**S1. Supplementary Materials and Methods**

- S1.1. Sensitivity Analysis: Greedy Age-Matching Algorithm

**S2. Supplementary Results**

- S2.1. Stability of Effects Across Motion Thresholds
- S2.2. Sensitivity Analysis in an Age-Matched Subsample
- S2.3. Duration-Matched Segment Analysis (*Despicable Me*)
- S2.4. Effects of Age and Sex
- S2.5. Age-Appropriate Tissue Probability Maps

**S3. Supplementary Discussion**

- S3.1. Theoretical Interpretation of Secondary Synchronization (PC2)
- S3.2. Developmental Maturation of Secondary Synchronization (PC2)

**S4. Supplementary References**

**S5. Supplementary Tables**

- Table S1. Demographic and Clinical Characteristics of the Sample at Different Head Motion Thresholds

**S6. Supplementary Figures**

- Figure S1. Age Distribution of the Samples
- Figure S2. Stability of Variance Partitioning Across Motion Thresholds during *The Present*
- Figure S3. Robustness of PC1 Effects and Variance Partitioning to Age Matching during *The Present*
- Figure S4. Variance Partitioned in Duration-Matched Segments during *Despicable Me*
- Figure S5. Variance Partitioning to Secondary Shared Responses (PC2) during *The Present*
- Figure S6. Developmental Maturation Effects (Age F-contrasts)
- Figure S7. Effect of Spatial Normalization Template on Extracted ROI Time Series and Principal Component Loadings
- Figure S8. Variance Partitioning Under Age-Matched Tissue Probability Maps

**S1. Supplementary Materials and Methods**

**S1.1. Sensitivity Analysis: Greedy Age-Matching Algorithm**

To assess whether the observed neural differences were driven by the age difference between the full ASD and TD cohorts, we performed a sensitivity analysis on an age-matched subsample. We employed a greedy nearest-neighbor matching algorithm to pair participants based on minimal age difference.

The algorithm proceeded in the following steps:

1. **Initialization:** The pool of eligible participants was defined as those meeting the primary motion threshold (Maximum FD < 2.4 mm/°).
2. **Iterative Nearest-Neighbor Selection:** The algorithm iterated through the Typically Developing (TD) cohort. For each TD participant:
   - We calculated the absolute age difference between the current TD participant and all currently available ASD participants.
   - The ASD participant with the minimum absolute age difference was identified as the "best match."
   - This pair was retained for the subsample, and the selected ASD participant was removed from the available pool to ensure unique, one-to-one matching (sampling without replacement).
3. **Verification:** This process continued until all feasible pairs were identified. The final matched sample (*N*_ASD_ = 123, *N*_TD_ = 123) was statistically verified to have no significant difference in mean age (*t* = 0.20, p > 0.8).

All primary PCA, GLM, and variance partitioning analyses were repeated on this subsample. The resulting statistical maps (PC1 and PC2 loadings) and variance decomposition plots were visually and statistically compared to the full-sample results to ensure consistency (see Supplementary Figure S3).

**S2. Supplementary Results**

**S2.1. Stability of Effects Across Motion Thresholds**

We adopted a multi-threshold strategy to address the inherent trade-off between data quality and sample representativeness. While stringent thresholds (e.g., 1.2 mm/°) minimize motion artifacts, they risk reducing sample size and introducing selection bias by excluding participants with more severe symptoms. Conversely, lenient thresholds (e.g., 4.8 mm/°) preserve representativeness but introduce higher noise levels. By examining effects across this continuum, we aimed to isolate robust neural signatures that remained consistent regardless of exclusion criteria.

**Robustness of Primary Synchronization (PC1)**: For the dominant shared response (PC1), effects associated with both categorical diagnosis and dimensional SRS-2 scores were highly robust. Key social brain regions, including the medial prefrontal cortex (mPFC), amygdala, and anterior temporal cortex, exhibited significant effects across all three thresholds (1.2, 2.4, and 4.8 mm/°; see Main Text Figure 2). Critically, the unique dimensional contribution of the left amygdala remained stable across thresholds. This consistency strongly suggests that these findings are not attributable to head motion (which would dominate at 4.8 mm/°) or the specific exclusion of severe ASD cases (which occurs at 1.2 mm/°).

We further verified this stability by visually inspecting unthresholded variance partitioning plots (Supplementary Figure S2). The spatial distribution of unique and shared variance was closely similar across thresholds, indicating that minor fluctuations in statistical significance (FDR correction) were likely due to sample size variations rather than instability in the underlying signal.

**Sensitivity of Secondary Synchronization (PC2):** Results for secondary synchronization (PC2) were more sensitive to threshold selection than those for PC1, though the pattern was not a simple gradient. The right posterior superior temporal sulcus (ROI 93) was significant at all three thresholds. The bilateral precuneus (ROIs 36 and 89) was significant at the intermediate and lenient thresholds (2.4 and 4.8 mm/°) but not at the strict threshold (1.2 mm/°), whereas the left temporoparietal junction (ROI 40) showed the opposite pattern, reaching significance at the strict and intermediate thresholds but not at the lenient one. The intermediate threshold was the only one at which all four regions were significant.

These two failure modes correspond to the two arms of the trade-off described above. The loss of the precuneus effects at the strict threshold is most plausibly a matter of power: the sample falls from 327 to 246 participants, a 25% reduction, and PC2 accounts for only about 5% of the variance. The loss of the temporoparietal effect at the lenient threshold is more consistent with added noise, and with the data-driven nature of PCA: because components are recomputed within each subsample, including a large number of high-motion participants can alter the component structure itself and shift variance away from PC2. That the two losses occur at opposite ends of the range supports treating 2.4 mm/° as the balance point between sample size and signal quality. This threshold was defined a priori as one voxel dimension (Main Text Section 2.4) and was used for all primary analyses, including PC1, rather than selected on the basis of the PC2 results.

**S2.2. Sensitivity Analysis in an Age-Matched Subsample**

Using the matching procedure described in Section S1.1, we selected a subset of participants (*N*_ASD_ = 123; *N*_TD_ = 123) balanced for age (*t* = 0.20, *p* > 0.8).

**Robustness of Primary Synchronization (PC1):** Despite the reduction in statistical power inherent to the smaller sample size (*N* = 246 vs. 327), the spatial topography of neural synchronization remained highly consistent with the full-sample analysis. At an uncorrected threshold of *p* < 0.05 (Supplementary Figure S3), we observed significant effects in all regions identified in the main analysis.

Crucially, the specific unique dimensional contribution of SRS-2 scores in the left amygdala remained statistically significant (*t* = -2.88, *p* = 0.0044) even after controlling for diagnosis. This indicates that the "dual-track" dissociation, where the amygdala tracks severity rather than category, is unlikely to be an artifact.

To validate these patterns independent of statistical thresholds, we compared the unthresholded variance partitioning maps (Supplementary Figure S3). The spatial distribution of unique and shared variance contributions across the cortex was closely similar to that of the full sample (Main Text Figure 3A), indicating that the underlying signal structure is stable and not driven by age confounds.

**Robustness of Secondary Synchronization (PC2):** All four regions showing unique dimensional associations in the full sample remained significant in the age-matched subsample at uncorrected *p* < 0.05: left precuneus (|*t*| = 2.76, *p* = 0.006), right precuneus (|*t*| = 3.18, *p* = 0.002), left temporoparietal junction (|*t*| = 2.23, *p* = 0.026), and right posterior superior temporal sulcus (|*t*| = 2.15, *p* = 0.033). Because the sign of PC2 loadings is mathematically arbitrary, absolute *t* values are reported.

**S2.3. Duration-Matched Segment Analysis (*Despicable Me*)**

*The Present* and *Despicable Me* differ in duration as well as in content (240 vs. 740 volumes after discarding the first 10 volumes of each run). Because the reliability of both the PCA synchronization estimates and the subsequent variance partitioning depends on the number of time points, differences between the two stimuli could in principle reflect duration rather than the stimuli themselves. We therefore repeated the *Despicable Me* analysis on three duration-matched segments of 240 volumes each: volumes 1–240, 251–490, and 501–740. Because 740 volumes do not divide evenly into three 240-volume blocks, 20 intervening volumes were not analyzed. Within each segment, PCA was performed separately on the segment’s time series, and the same group-level general linear model and variance partitioning were applied, using the same participants as the full-run analysis (motion threshold 2.4 mm/°, n = 242). Results are shown in Figure S4.

The segment analyses did not reproduce the full-run partition. Spatial correlations between each segment’s three variance components and those of the full run were modest to moderate (shared: r = 0.51, 0.09, 0.68; SRS-unique: r = 0.09, 0.33, 0.72; diagnosis-unique: r = 0.35, 0.38, 0.69 for the first, second, and third segment respectively). These values are, however, inflated: each segment contributes roughly one third of the data entering the full-run estimate. The non-circular comparison is between segments, and there the correlations were close to zero (shared: r = −0.13, 0.27, −0.28; SRS-unique: r = 0.00, −0.06, 0.07; diagnosis-unique: r = −0.02, 0.18, 0.12 for segment 1 vs. 2, 1 vs. 3, and 2 vs. 3). Variance partitioning estimated within a single 240-volume window of *Despicable Me* is therefore not stable across windows, and we do not interpret the segment-level maps substantively.

Two considerations bear on this instability. First, it is not attributable to duration alone, because *The Present* comprises the same number of volumes and yielded a partition that was reproducible across head motion thresholds (Supplementary Section S2.1) and across tissue probability maps (Section S2.5). Second, the two stimuli differ in narrative structure in a way that duration matching does not address. *The Present* is a complete short film with a single story arc, designed to be viewed in its entirety, so its 240 volumes constitute a whole narrative. *Despicable Me*, by contrast, is a 10-minute excerpt from a feature-length film and contains several distinct story segments within it; an arbitrary 240-volume window therefore samples a fragment of a longer narrative rather than a self-contained one, and different windows sample qualitatively different material. This is consistent with reports that the magnitude and spatial distribution of inter-subject synchronization vary considerably across film stimuli (Leipold et al., 2026), and it indicates that duration, narrative completeness, and content are not separable in the present design. We accordingly retain the full-run *Despicable Me* analysis as the comparison of interest in the main text and treat stimulus duration as one of several uncontrolled differences between the two runs rather than as an isolable factor.

**S2.4. Effects of Age and Sex**

We modeled non-linear developmental trajectories using both log-transformed age and its square. For the dominant shared response (PC1), age effects exhibited a distinct spatial topography compared to clinical effects, clustering primarily within the dorsal attention network and somatomotor regions (Supplementary Figure S6). Notably, the social-motivational hubs identified in the main analysis, specifically the medial prefrontal cortex, caudate, and amygdala, did not exhibit significant age-related modulation, supporting a functional dissociation between normative maturation and the specific neural signatures of ASD.

In contrast, the secondary response pattern (PC2) showed more widespread developmental effects, extending into lateral occipital and temporal cortices. The precuneus, temporoparietal junction, and posterior superior temporal sulcus regions that showed dimensional associations with SRS-2 scores also exhibited significant age-related synchronization. However, because age terms were included as covariates in all primary analyses, the reported SRS-2 effects in these regions reflect unique variance associated with social impairment above and beyond normative developmental change. No significant main effects of sex were observed.

**S2.5. Age-Appropriate Tissue Probability Maps**

To evaluate whether the findings depend on the use of adult-derived tissue priors, we repeated spatial normalization using tissue probability maps matched to the age distribution of our sample (Wilke et al., 2008), and re-extracted ROI time series and principal component loadings. Agreement between pipelines was quantified as the correlation across time points between each participant’s ROI time series under the two pipelines, and as the correlation across participants between PC loadings under the two pipelines.

Time series agreement. Agreement was high and essentially identical across stimuli. For *The Present* (n = 327 × 114 ROIs), correlations had a mean of 0.995 and a median of 0.997, with 97.1% of values ≥ 0.98. For *Despicable Me* (n = 242 × 114), the mean was 0.995 and the median 0.997, with 97.8% ≥ 0.98. All 114 ROI medians were ≥ 0.98 in both runs (Supplementary Figure S7A, B). The lowest agreement occurred in small structures adjacent to tissue boundaries — pallidum, parahippocampal gyrus, and amygdala — with medians of 0.984–0.987. For the left amygdala, the region carrying the unique dimensional effect, median agreement was 0.987 (*The Present*) and 0.988 (*Despicable Me*).

Values below r = 0.95 were rare (90 of 37,278 in *The Present*; 77 of 27,588 in *Despicable Me*) and were attributable to a small number of individual participants rather than to particular regions (73.3% arising from five participants in *The Present*). Visual inspection indicated tissue misclassification in the age-matched segmentations of two of these participants, aged 13 and 14 years — isolated segmentation failures unrelated to participant age, since the age-matched priors are well suited to this range.

Component loadings. PC1 loadings were highly consistent across all regions (*The Present*: median 0.996, range 0.978–0.999, left amygdala r = 0.982; *Despicable Me*: median 0.997, range 0.968–1.000, left amygdala r = 0.990). PC2 loadings were likewise consistent in most regions (*The Present*: median |r| = 0.995; 6 of 114 regions with |r| < 0.95; *Despicable Me*: median |r| = 0.996; 7 of 114 regions with |r| < 0.95), with lower agreement where PC2 is weakly expressed, consistent with its smaller share of variance. The regions underlying the reported PC2 effect (ROIs 36, 40, 89, and 93) showed high agreement (|*r*| = 0.997, 0.995, 0.992, and 0.994; Supplementary Figure S7C, D).

Repeating the group-level analyses under the age-matched pipeline yielded similar patterns of dimensional and categorical association. The unique dimensional association with SRS-2 scores in the left amygdala remained significant under the same masked FDR correction (t = −3.24, p = 0.0013), and the shared diagnosis/SRS-2 contributions in the mPFC and caudate retained the same pattern (Supplementary Figure S8A). Despicable Me partitioning was likewise unchanged (Supplementary Figure S8B).

Given that the age-matched priors produced near-identical time series while introducing a small number of segmentation failures, we retained the standard tissue probability maps for the primary analyses and report this as a sensitivity analysis.

**S3. Supplementary Discussion**

**S3.1. Theoretical Interpretation of Secondary Synchronization (PC2)**

Our previous work indicates that secondary principal components in movie fMRI often capture temporal offsets relative to the primary shared response (Di and Biswal, 2022). Treating PC2 as temporal structure rather than residual noise is further supported by evidence that BOLD response profiles during naturalistic viewing depart substantially from the canonical hemodynamic response (Di et al., 2026).

The regions carrying the PC2 effect, the bilateral precuneus, the left temporoparietal junction, and the right posterior superior temporal sulcus, are adjacent to, but distinct from, the PC1 hubs in the posterior cingulate and anterior temporal cortex. In the Schaefer parcellation the precuneus parcels belong to the control network and the parietal and temporal parcels to the default mode network, the latter two being canonical nodes of the mentalizing network. This arrangement suggests a functional gradient rather than a duplication: the posterior cingulate and medial prefrontal regions that dominate PC1 track the shared narrative directly, whereas neighboring posterior-medial and lateral association cortex contributes variance that is offset in time from that signal.

On this reading, the unique association between SRS-2 scores and PC2 loadings reflects altered timing in the higher-order integration of narrative content rather than a difference in how strongly the narrative is tracked. Because the sign of PC2 loadings is mathematically arbitrary, our data do not indicate whether this constitutes a lead or a lag. An account of this kind is compatible with proposals that autism involves altered precision in predictive processing, which would be expected to affect the rate at which internal models are updated during a continuously unfolding narrative (Van de Cruys et al., 2014).

**S3.2. Developmental Maturation of Secondary Synchronization (PC2)**

Age effects on the secondary component were more widespread than those on the primary component (Section S2.4), consistent with our earlier report in an independent sample, in which developmental effects were likewise stronger for PC2 than for PC1 (Di and Biswal, 2022). Because PC2 captures response structure that is temporally offset from the dominant shared signal, this pattern suggests that the temporal organization of narrative processing continues to change across childhood and adolescence, whereas the strength of narrative tracking is comparatively stable across this range.

This developmental sensitivity bears on the interpretation of the PC2 findings reported here. The precuneus, temporoparietal junction, and posterior superior temporal sulcus showed both age effects and dimensional associations with SRS-2 scores (Section S2.4), raising the possibility that the two reflect a single underlying gradient, with higher symptom severity corresponding to a less mature temporal profile. The group-level models address this directly: log(age) and [log(age)]² were included as covariates throughout, so the reported SRS-2 associations describe variance not accounted for by normative developmental change. The association in these regions is therefore not reducible to delayed maturation, although both influences act on the same component and the present design cannot establish that they are mechanistically distinct.

**Supplementary References:**

Di, X., Biswal, B.B., 2022. Principal component analysis reveals multiple consistent responses to naturalistic stimuli in children and adults. Human Brain Mapping 43, 3332–3345. https://doi.org/10.1002/hbm.25568

Di, X., Hanna, G.B., Biswal, B.B., 2026. Beyond the Canonical HRF: Flexible Temporal Modeling Reveals Unconstrained BOLD Profiles During Naturalistic Viewing. Imaging Neuroscience. https://doi.org/10.1162/IMAG.a.1331

Leipold, S., Rao, R.R., Schoffelen, J.-M., Bögels, S., Toni, I., 2026. Inter-subject correlations and their behavioral associations vary across movies: Implications for generalizability. Imaging Neuroscience. https://doi.org/10.1162/IMAG.a.1400

Van de Cruys, S., Evers, K., Van der Hallen, R., Van Eylen, L., Boets, B., de-Wit, L., Wagemans, J., 2014. Precise minds in uncertain worlds: predictive coding in autism. Psychol Rev 121, 649–675. https://doi.org/10.1037/a0037665

Wilke, M., Holland, S.K., Altaye, M., Gaser, C., 2008. Template-O-Matic: a toolbox for creating customized pediatric templates. Neuroimage 41, 903–913. https://doi.org/10.1016/j.neuroimage.2008.02.056

**Supplementary Table S1.** Participants characteristics for the two groups, and at different head motion threshold levels for *The Present* and *Despicable Me*. * indicates statistical significance at p < 0.05 between ASD and TD groups for sex, age, Social Responsiveness Scale-2 (SRS-2), and mean framewise displacement (FD).

| Video | | *The Present* | | | | | | *Despicable Me* | | | | | |
| --- | --- | --- | --- | --- | --- | --- | --- | --- | --- | --- | --- | --- | --- |
| Max FD threshold | | 4.8 | | 2.4 | | 1.2 | | 4.8 | | 2.4 | | 1.2 | |
| Group | | ASD | TD | ASD | TD | ASD | TD | ASD | TD | ASD | TD | ASD | TD |
| n | All | 262 | 166 | 204 | 123 | 150 | 96 | 202 | 141 | 141 | 101 | 90 | 56 |
|  | Female | 45 | 67 | 36 | 53 | 28 | 42 | 35 | 59 | 29 | 41 | 21 | 23 |
|  | Χ^2^, p | 28.27 | * <0.01 | 25.07 | * <0.01 | 18.09 | * <0.01 | 25.09 | * <0.01 | 11.48 | * <0.01 | 5.16 | * 0.02 |
| Age | Mean | 11.16 | 10.02 | 11.51 | 10.29 | 11.77 | 10.65 | 11.63 | 10.26 | 11.93 | 10.49 | 12.23 | 11.42 |
|  | STD | 3.51 | 3.27 | 3.57 | 3.50 | 3.67 | 3.60 | 3.55 | 3.34 | 3.72 | 3.65 | 3.90 | 3.83 |
|  | t, p | 3.36 | * <0.01 | 3.00 | * <0.01 | 2.35 | 0.02 | 3.58 | * <0.01 | 2.98 | * <0.01 | 1.24 | 0.22 |
| SRS | Mean | 68.60 | 48.59 | 68.92 | 48.98 | 69.32 | 48.91 | 68.76 | 49.15 | 68.53 | 50.03 | 69.08 | 49.41 |
|  | STD | 11.20 | 6.95 | 10.90 | 7.35 | 10.49 | 7.52 | 10.98 | 7.21 | 10.76 | 7.57 | 11.12 | 6.96 |
|  | t, p | 20.63 | * <0.01 | 17.97 | * <0.01 | 16.53 | * <0.01 | 18.59 | * <0.01 | 14.84 | * <0.01 | 11.86 | * <0.01 |
| Mean FD  translation | Mean | 0.13 | 0.12 | 0.11 | 0.10 | 0.10 | 0.09 | 0.12 | 0.12 | 0.10 | 0.11 | 0.09 | 0.09 |
|  | STD | 0.06 | 0.05 | 0.05 | 0.04 | 0.04 | 0.03 | 0.05 | 0.06 | 0.04 | 0.04 | 0.03 | 0.03 |
|  | t, p | 1.83 | 0.07 | 1.60 | 0.11 | 1.75 | 0.08 | -0.27 | 0.78 | -0.79 | 0.43 | -0.60 | 0.55 |
| Mean FD  rotation | Mean | 0.08 | 0.08 | 0.06 | 0.06 | 0.05 | 0.04 | 0.07 | 0.08 | 0.05 | 0.06 | 0.04 | 0.04 |
|  | STD | 0.06 | 0.05 | 0.03 | 0.03 | 0.02 | 0.01 | 0.05 | 0.05 | 0.02 | 0.04 | 0.01 | 0.02 |
|  | t, p | 0.81 | 0.42 | 0.73 | 0.47 | 0.91 | 0.36 | -0.74 | 0.46 | -1.51 | 0.13 | 0.33 | 0.75 |

**Supplementary Figure S1. Age distributions of the samples.** Histograms at the primary head motion threshold of 2.4 mm/° for *The Present* (**A**; *n* = 327) and *Despicable Me* (**B**; *n* = 242).

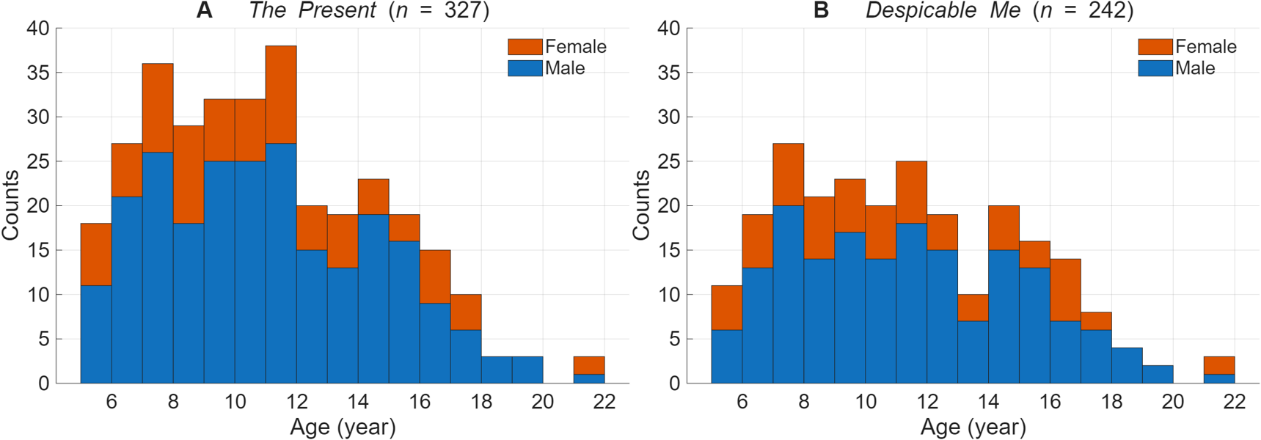

**Supplementary Figure S2.** **Variance partitioning of dimensional and categorical contributions to primary neural synchronization during *The Present*.** The figure illustrates the decomposition of variance in synchronization (PC1) into unique contributions from Social Responsiveness Scale-2 (SRS-2) Total T-scores, unique contributions from clinical diagnosis (ASD vs. TD), and their shared variance across 114 regions of interest. Data are presented for three head motion thresholds of 1.2, 2.4, and 4.8 mm/°.

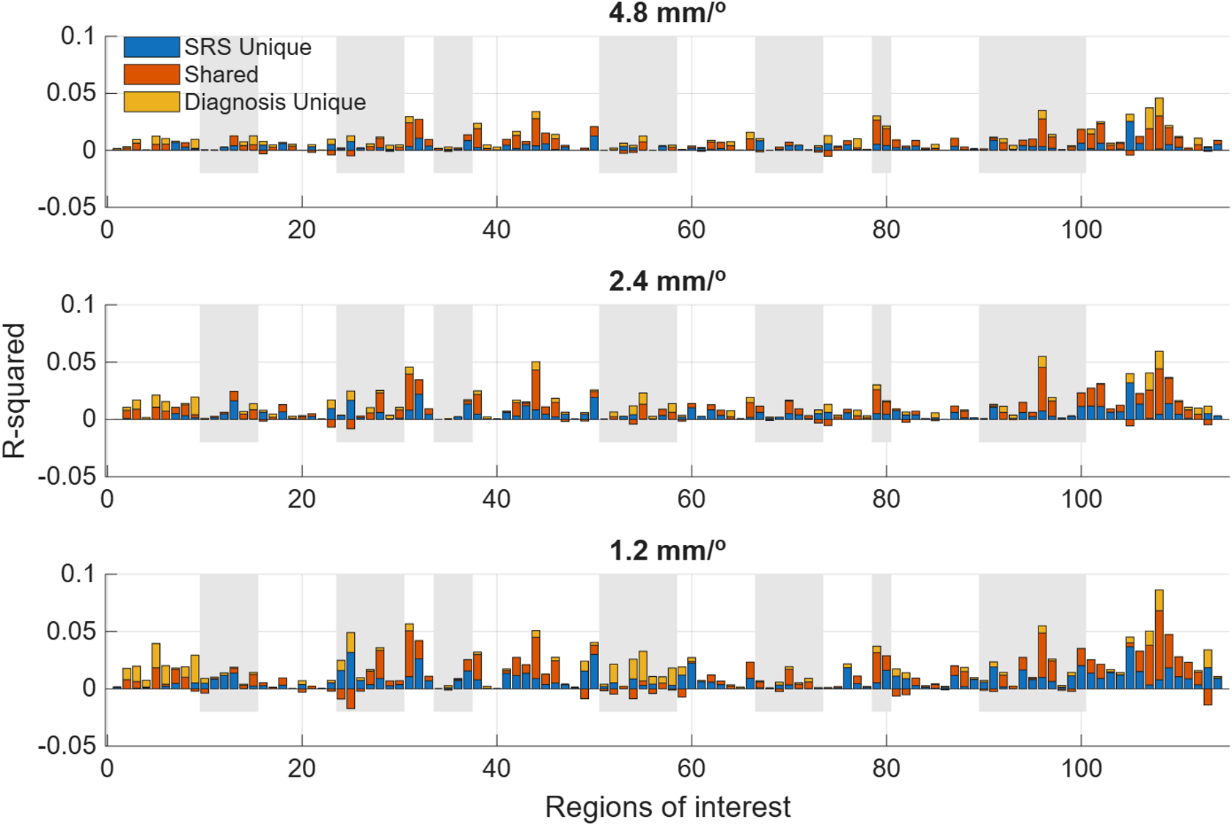

**Supplementary Figure S3. Robustness of dimensional and categorical effects to age matching during *The Present*.** Sensitivity analyses of primary neural synchronization (PC1) performed in a age-matched subsample (*N*_ASD_ = 123, *N*_TD_ = 123) at the primary head motion threshold of 2.4 mm/°. **(A)** Dimensional association between continuous symptom severity (SRS-2 Total T-scores) and PC1 loadings. **(B)** Categorical group differences in PC1 loadings (TD > ASD). Statistical maps in (A) and (B) are thresholded at uncorrected *p* < 0.05. **(C)** Decomposition of variance in PC1 across the 114 regions of interest into the unique contribution of SRS-2 Total T-scores, the unique contribution of categorical diagnosis, and the variance shared between them. Negative values reflect suppression effects rather than negative variance explained. Across all three panels, the spatial topography closely resembles the corresponding full-sample results (Main Text Figures 2 and 3), indicating that the reported effects are unlikely to be attributable to age differences between groups.

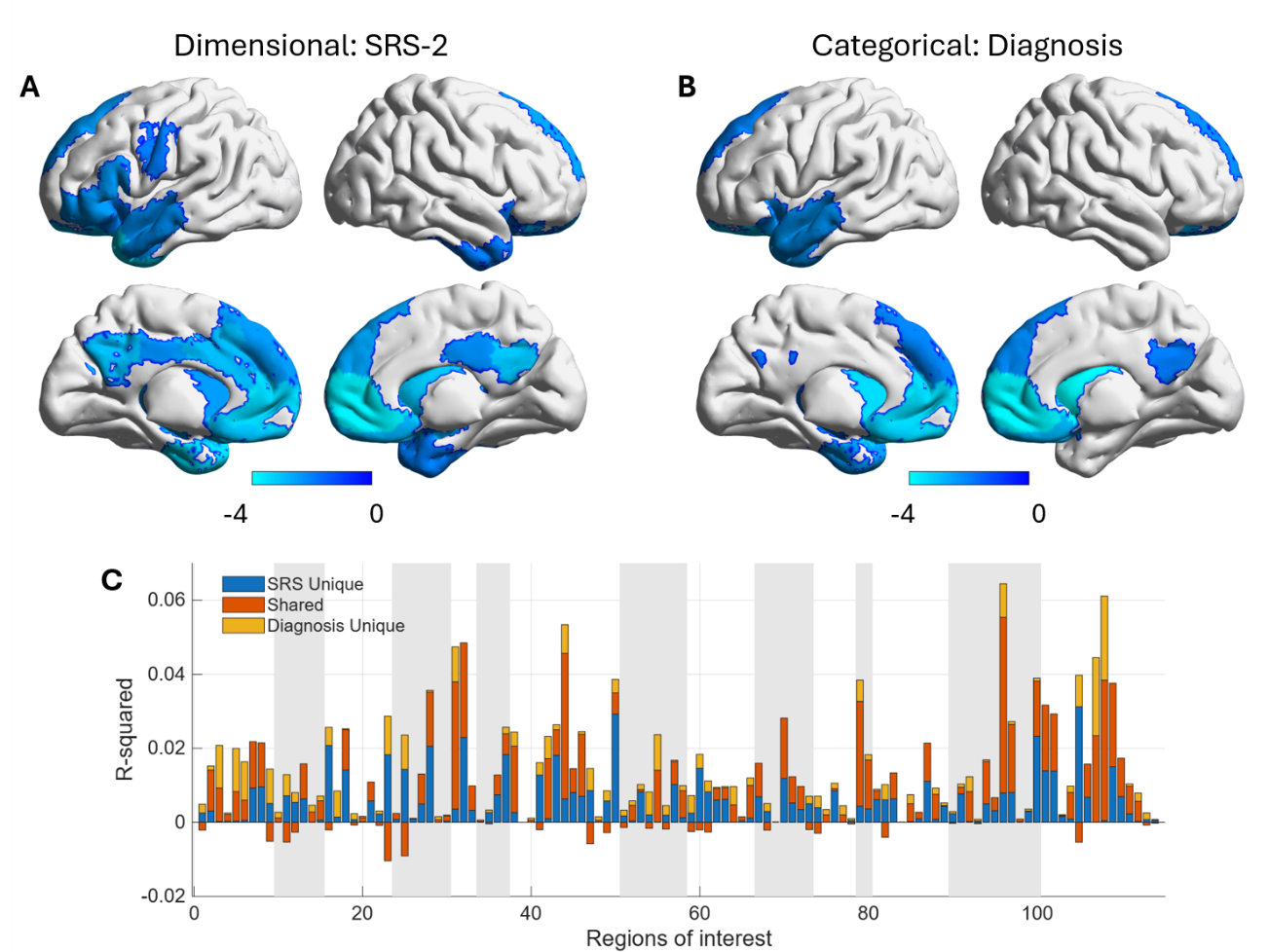

**Supplementary Figure S4.** **Variance partitioning of dimensional and categorical contributions to neural synchronization in duration-matched segments of *Despicable Me*.** Each panel decomposes the variance in regional synchronization (PC1) into the unique contribution of Social Responsiveness Scale-2 (SRS-2) Total T-scores, the unique contribution of clinical diagnosis (ASD vs. TD), and the variance shared between the two predictors, across 114 regions of interest. Panels correspond to three non-overlapping 240-volume segments, matched in duration to *The Present*: (A) volumes 1–240, (B) volumes 251–490, and (C) volumes 501–740; the 20 intervening volumes were not analyzed. Principal component analysis was performed separately within each segment, and the group-level model was identical to that of the full-run analysis. Regions are ordered as in Main Text Figure 3. Negative shared values indicate suppression, where the two predictors jointly account for more variance than the sum of their separate contributions. Data are shown for the primary head motion threshold (2.4 mm/°; n = 242).

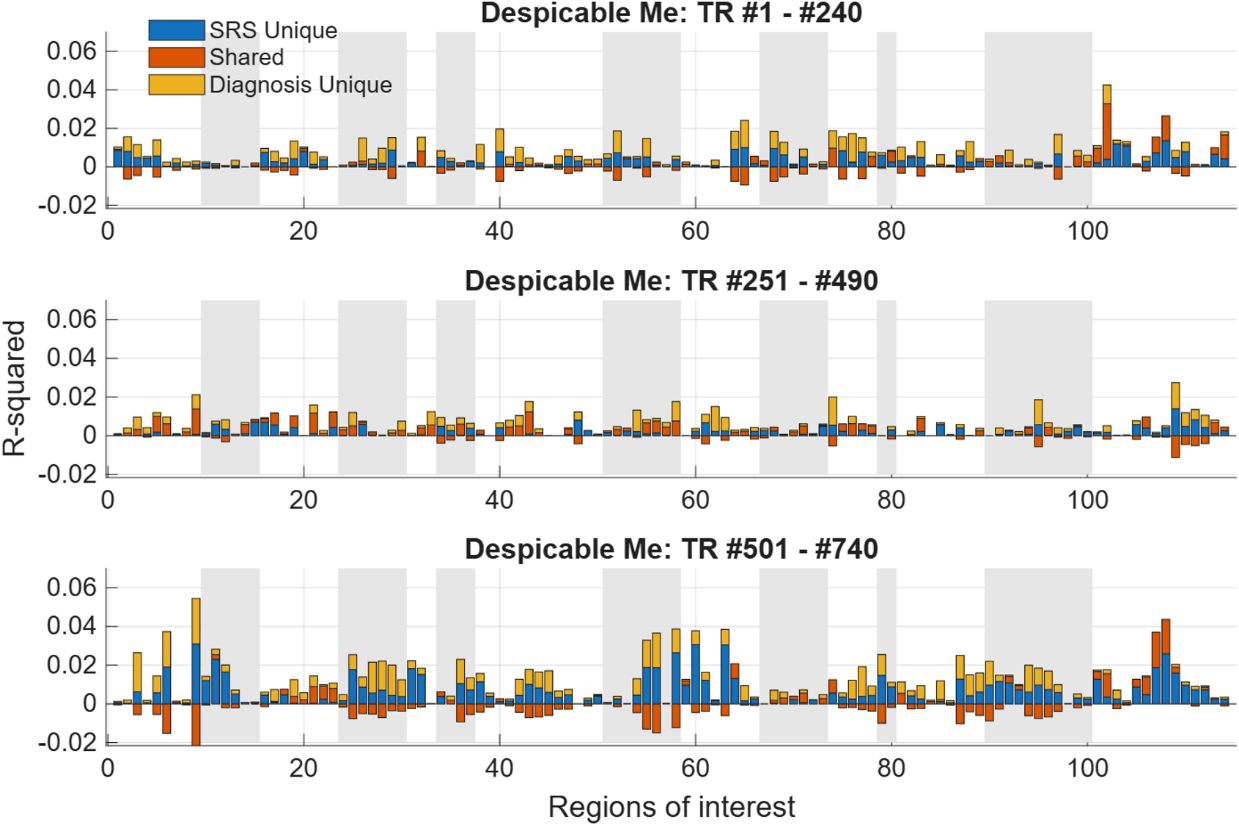

**Supplementary Figure S5.** **Variance partitioning of dimensional and categorical contributions to secondary shared response patterns (PC2) during *The Present*.** This figure decomposes the variance in secondary shared regional response (PC2) into the unique contribution of Social Responsiveness Scale-2 (SRS-2) Total T-scores, the unique contribution of clinical diagnosis (ASD vs. TD), and the variance shared between the two predictors, across 114 regions of interest. Negative shared values indicate suppression, where the two predictors jointly account for more variance than the sum of their separate contributions. Data are shown for the primary head motion threshold (2.4 mm/°; n = 327).

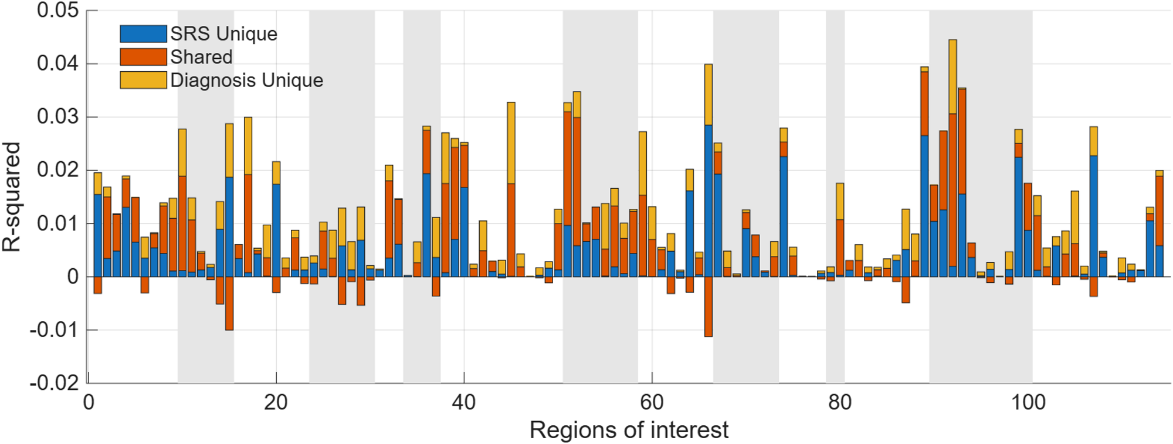

**Supplementary Figure S6. Developmental maturation effects on neural synchronization during *The Present*.** The top row illustrates age-related effects on the dominant shared response (PC1), while the bottom row displays effects on the secondary response pattern (PC2). Age effects were evaluated using an *F*-contrast incorporating both log(age) and [log(age)]^2^ terms to capture potential non-linear developmental trajectories. Left panels present *F*-statistics at the primary head motion threshold of 2.4 mm/°. Right panels show the consistency of these maturational effects across three motion thresholds (1.2, 2.4, and 4.8 mm/°). The color scale represents the number of thresholds at which the effect reached significance, with values of 1, 2, and 3 denoting consistency across one, two, or all three thresholds, respectively. All statistical maps were thresholded at a False Discovery Rate (FDR) of *p* < 0.05 across 114 regions of interest.

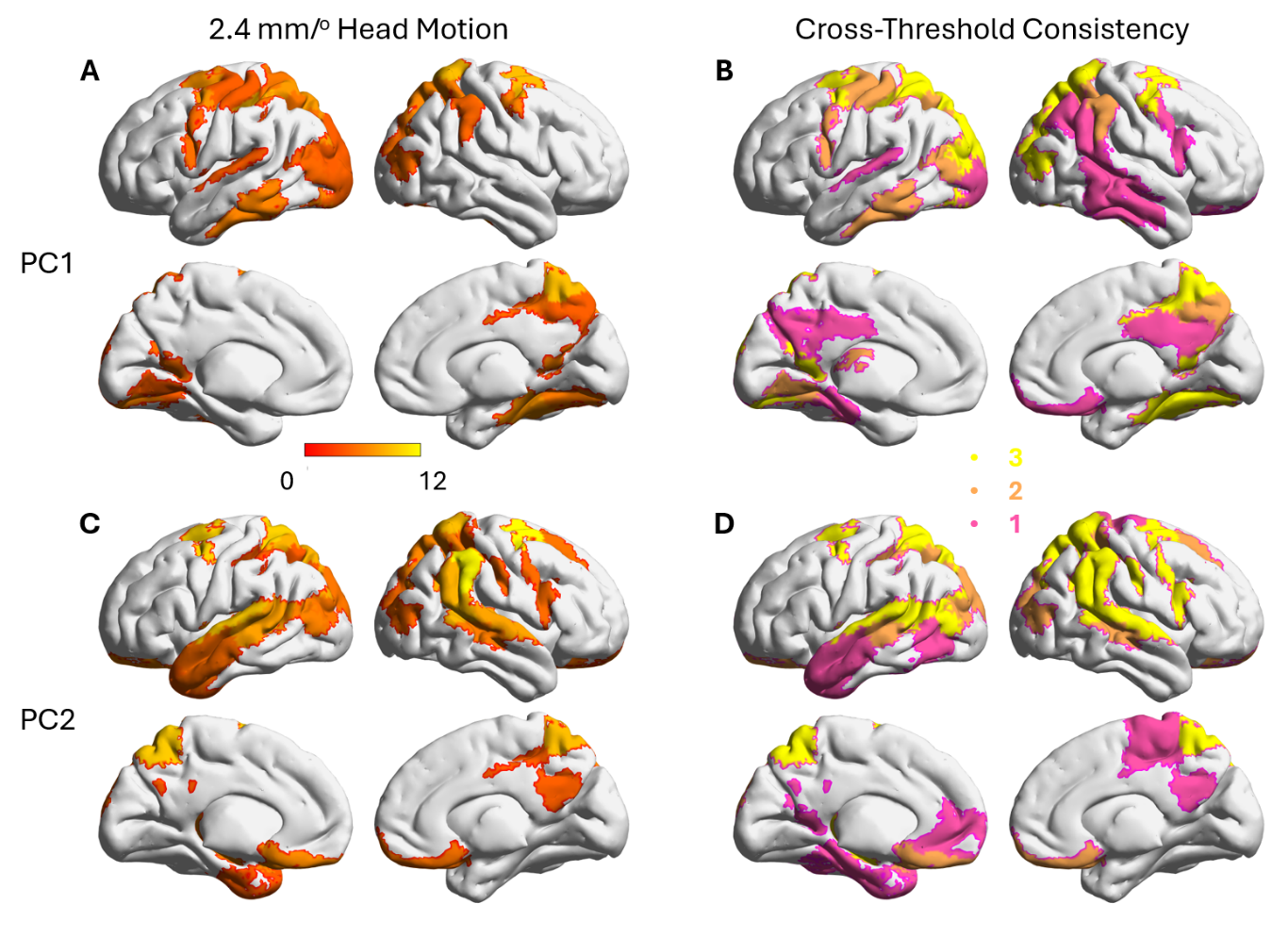

**Supplementary Figure S7. Effect of spatial normalization template on extracted ROI time series and principal component loadings.** Comparison between data processed using SPM12’s default tissue probability maps and age-matched tissue probability maps generated with the Template-O-Matic toolbox, at the primary head motion threshold of 2.4 mm/°. **(A, B)** Median correlation across participants between ROI time series derived under the two pipelines, for each of the 114 regions of interest, during *The Present* (A; *n* = 327) and *Despicable Me* (B; *n* = 242). Error bars denote the 5th and 95th percentiles across participants. **(C, D)** Correlations across participants between the first (PC1) and second (PC2) principal component loadings derived under the two pipelines, for each region. Because the sign of PC loadings is mathematically arbitrary, correlations are shown as absolute values. Shaded bands indicate network boundaries; regions 101–114 are subcortical.

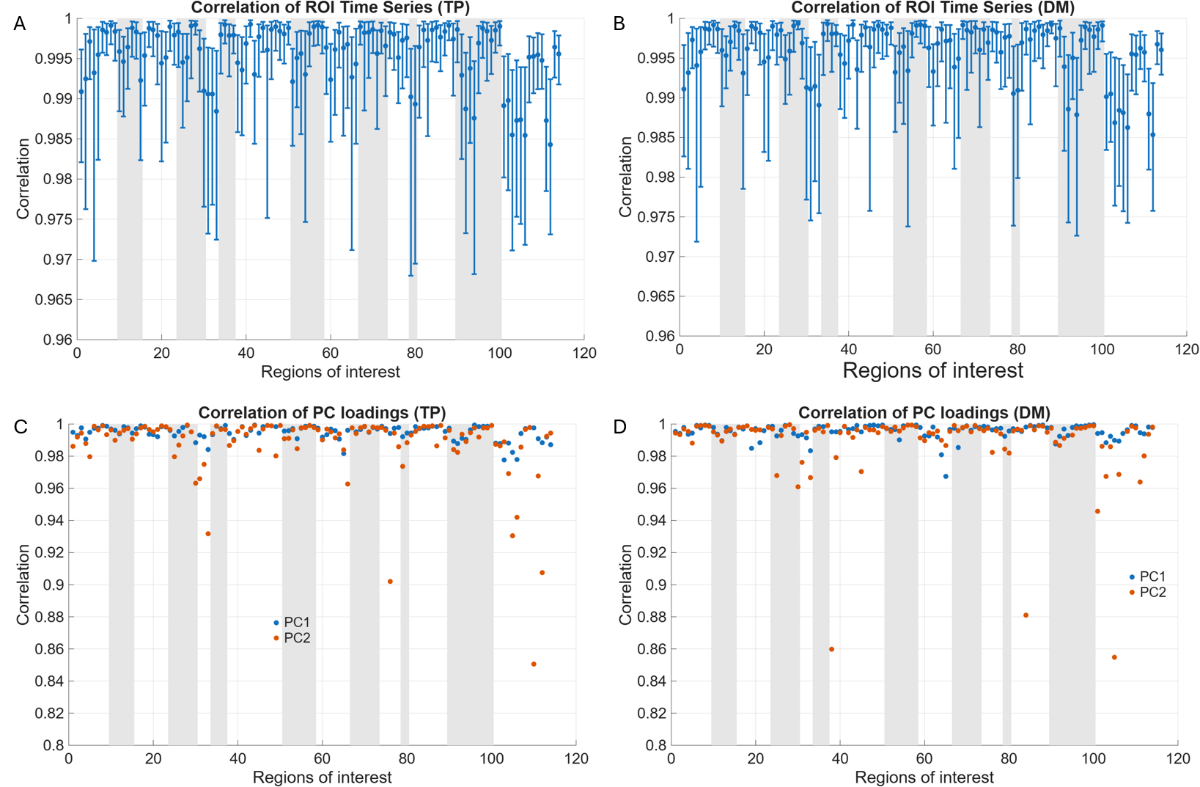

**Supplementary Figure S8. Variance partitioning under age-matched tissue probability maps during *The Present* (A) and *Despicable Me* (B).** Decomposition of variance in primary synchronization (PC1) into the unique contribution of SRS-2 Total T-scores, the unique contribution of categorical diagnosis, and the variance shared between them, across 114 regions of interest, at the primary head motion threshold of 2.4 mm/° (*The Present*, *n* = 327; *Despicable Me*, *n* = 242). Data were processed using age-matched tissue probability maps generated with the Template-O-Matic toolbox. Negative values reflect suppression effects rather than negative variance explained. The spatial patterns closely resemble those obtained with the standard tissue probability maps (Main Text Figure 3A, B).

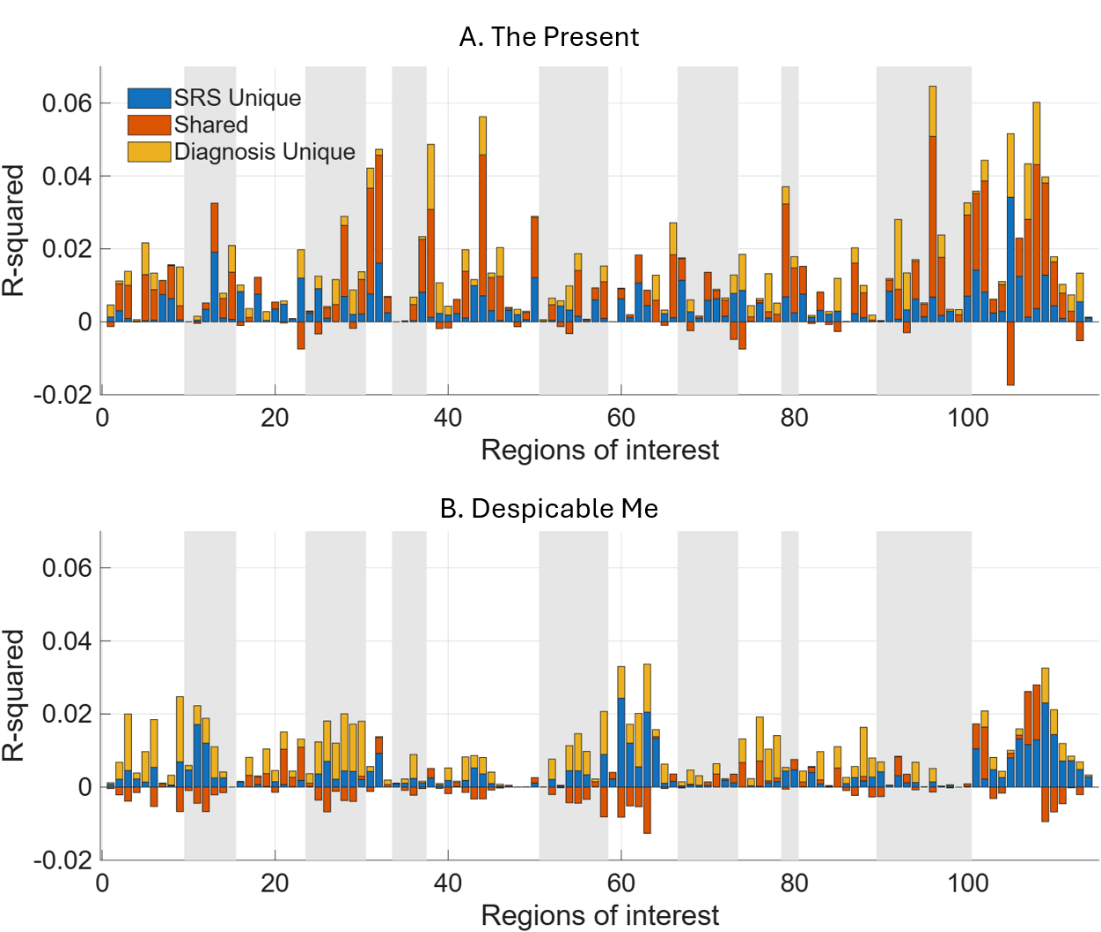
